# Multiformity of extracellular microelectrode recordings from Aδ neurons in the dorsal root ganglia: A computational modeling study

**DOI:** 10.1101/2023.10.17.562569

**Authors:** Lauren R. Madden, Robert D. Graham, Scott F. Lempka, Tim M. Bruns

## Abstract

Microelectrodes serve as a fundamental tool in electrophysiology research throughout the nervous system, providing a means of exploring neural function with a high resolution of neural firing information. We constructed a hybrid computational model using the finite element method and multi-compartment cable models to explore factors that contribute to extracellular voltage waveforms that are produced by sensory pseudounipolar neurons — specifically, smaller A-type neurons — and that are recorded by microelectrodes in dorsal root ganglia. The finite element method model included a dorsal root ganglion, surrounding tissues, and a planar microelectrode array. We built a multi-compartment neuron model with multiple trajectories of the glomerular initial segment found in many A-type sensory neurons. Our model replicated both the somatic intracellular voltage profile of Aδ low-threshold mechanoreceptor neurons and the unique extracellular voltage waveform shapes that are observed in experimental settings. Results from this model indicated that tortuous glomerular initial segment geometries can introduce distinct multiphasic properties into a neuron’s recorded waveform. Our model also demonstrated how recording location relative to specific microanatomical components of these neurons, and recording distance from these components, can contribute to additional changes in the multiphasic characteristics and peak-to-peak voltage amplitude of the waveform. This knowledge may provide context for research employing microelectrode recordings of pseudounipolar neurons in sensory ganglia, including functional mapping and closed-loop neuromodulation. Further, our simulations gave insight into the neurophysiology of pseudounipolar neurons by demonstrating how the glomerular initial segment aids in increasing the resistance of the stem axon and mitigating rebounding somatic action potentials.

## Introduction

Neural electrophysiology techniques have long been supported by microelectrode technology development. Neural interfacing with microelectrodes in the brain is commonly used to explore the neurophysiology of different cortical areas due to the ability of these electrodes to record spikes of single neural units [1]–[3]. Ascribable to their neuron-level specificity, microelectrodes have been used to record single-unit activity correlated with movement intent and to stimulate sensory cortical areas to evoke cutaneous sensation [4]. Neural electrophysiology research employing penetrating microelectrodes has also played a similar role in answering questions about the peripheral nervous system and the numerous neuromodulation therapies that target its structures. Microelectrode recordings have contributed to the understanding of peripheral neurophysiology [5]–[8] and also provided high-resolution signal inputs for exploring machine-learning techniques driving closed-loop stimulation methods [9], [10].

One common recording target for peripheral neurophysiology research, dorsal root ganglia (DRG), are clusters of neural cell bodies adjacent to, but outside of, the spinal cord. With bilateral pairs at each spinal level, DRG contain the cell bodies of neurons that innervate individual dermatomes and their respective organs, making them a discrete target for recording both somatic and visceral sensory information. Many peripheral electrophysiology studies have recorded single-unit activity at different peripheral locations, including the DRG, in response to various types of sensory stimuli [11]–[14].

Notably, unit waveforms recorded in the DRG with microelectrodes often take on shapes that are quite distinct from those seen in cortical and peripheral axon targets [14] (*Figure 1A*). Occasionally, waveforms recorded in the DRG have multiphasic, W-shaped signals, suggesting differences in the recording conditions, electrophysiology, or morphology of the neurons that give rise to them (*Figure 1B*). There is a lack of understanding of the factors that contribute to the extracellular waveforms of DRG neurons – factors that are neurophysiological, microanatomical, or dependent on experimental variations. This knowledge may inform how microelectrode interfacing with DRG neurons can be improved with respect to signal interpretation. It may also provide context for how penetrating microelectrode designs can be tailored to capture different types of neural signals. Further, an improved signal interpretation, rooted in an understanding of DRG neuron electrophysiology and morphology, may aid development of automated spike sorting algorithms for closed-loop peripheral neuromodulation [10].

**Figure 1:**
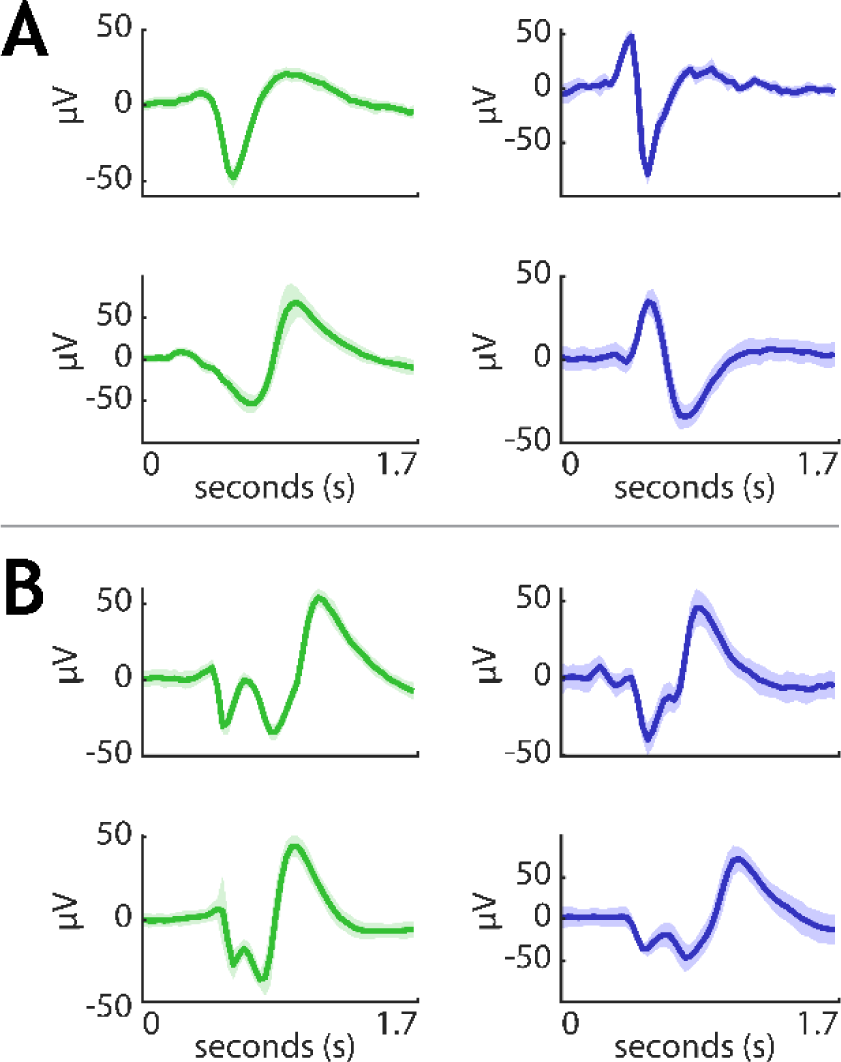
Example in vivo extracellular DRG waveforms. Bold lines show mean waveforms, and shading shows respective standard deviations. All waveforms were recorded from a feline sacral DRG. Green waveforms in the left columns were recorded with a Utah array, and blue waveforms in the right columns were recorded with the microelectrode array described in Sperry et al. 2021 [14]. (A) Typical waveform shapes that are often found in extracellular DRG neuron recordings. (B) Examples of W-shaped DRG waveforms that are occasionally observed.

Unique to sensory ganglia, neurons in the DRG have a pseudounipolar morphology. They have a stem axon protruding from the soma that splits at a T-junction into a peripheral branch, that projects distally, and a central branch, that enters the spinal cord. Some A-type neurons in the DRG have a long, tortuous initial segment of nonmyelinated axon near the soma known as a glomerulus [15]. Glomeruli are not unique to the DRG, as they have also been found in other sensory ganglia neurons, such as the trigeminal and vagal ganglia [16]. The function and evolutionary advantage of glomeruli are not well understood. It has also yet to be determined whether auto-ephaptic interactions play a role in their electrophysiology. Ephaptic coupling, the phenomenon that the electric field generated by a firing neuron can influence the firing properties of adjacent neurons [17], [18], has been demonstrated *in vitro*, *in silico*, and in both the central and peripheral nervous systems [19]–[22]. The impact of the extracellular electric field produced by a firing axon on other non-firing areas of the same axon (i.e., auto-ephaptic coupling) has yet to be investigated.

However, studying the electrophysiological and geometrical properties of glomeruli that affect recorded waveforms would be challenging to investigate *in vivo*. Instead, a computational recording model of the DRG and their uniquely structured neurons would provide a practical method of investigating glomeruli function and the effects of their tortuosity on recorded waveforms. Computational modeling of microelectrode recordings has informed electrophysiology research within the brain [23]–[25] and peripheral nerves [26], yet the factors that contribute to DRG microelectrode recordings have not been explored in this manner.

Here, we constructed a hybrid computational model to explore the factors that influence extracellular waveforms recorded in a feline DRG with a microelectrode. The first component was a volumetric model of a feline sacral level DRG and a single-shank, multi-site microelectrode array with which we performed finite element analysis on the voltage distribution around the contacts within gross tissue volumes. The second component was a multi-compartment double cable model of an Aδ neuron, complete with a peripheral branch, central branch, stem axon, glomerulus, and soma. This neuron model, and geometric variations of it, demonstrated how the location of the recording contact and structure of the neuron can influence extracellular voltage recordings. We also implemented a source summation method to investigate the effect of auto-ephaptic coupling within the glomerulus of the neuron as a mechanism for preventing the back-propagation of action potentials towards the T-junction of the neuron.

## Methods

We constructed a hybrid model to simulate extracellular recordings of DRG neurons, based on prior *in vivo* observations [14], [27]. We constructed a finite element method (FEM) model, built in COMSOL (COMSOL Inc., USA) [28], to model a sacral feline DRG, surrounding tissues, and a planar multielectrode array. We built a multi-compartment double cable model in the NEURON simulation environment [29] to simulate the morphology and membrane dynamics of an Aδ low-threshold mechanoreceptor (Aδ-LTMR). Aδ-LTMR neurons, a classification of smaller A-type sensory fibers, primarily transmit nonpainful touch and bladder state information and are a primary focus of our research [30], [31], [14]. We coupled the FEM model with the NEURON model, and we varied several components of this hybrid model to explore how these factors influence extracellular neural recordings. The model code can be found online at https://doi.org/10.5281/zenodo.10014936.

### FEM Model

The volume conductor FEM model included a sacral DRG, the respective dorsal root, a segment of the respective spinal nerve, a dural sheath around the neural tissues, a gross spinal cord, intraforaminal (and epidural) space, a gross model of vertebral bone, and general thoracic tissue (*Figure 2A-B*). We obtained the feline tissue dimensions (*Table 1*) and conductivity parameters (*Table 2*) from the literature and unpublished surgical images from prior *in vivo* experiments. We also included a model of a single-shank, multi-site, microelectrode array [14], [27] in the FEM model and placed it at the center of the DRG protruding dorsally to simulate *in vivo* experimental conditions [14] (*Figure 2C*).

**Figure 2:**
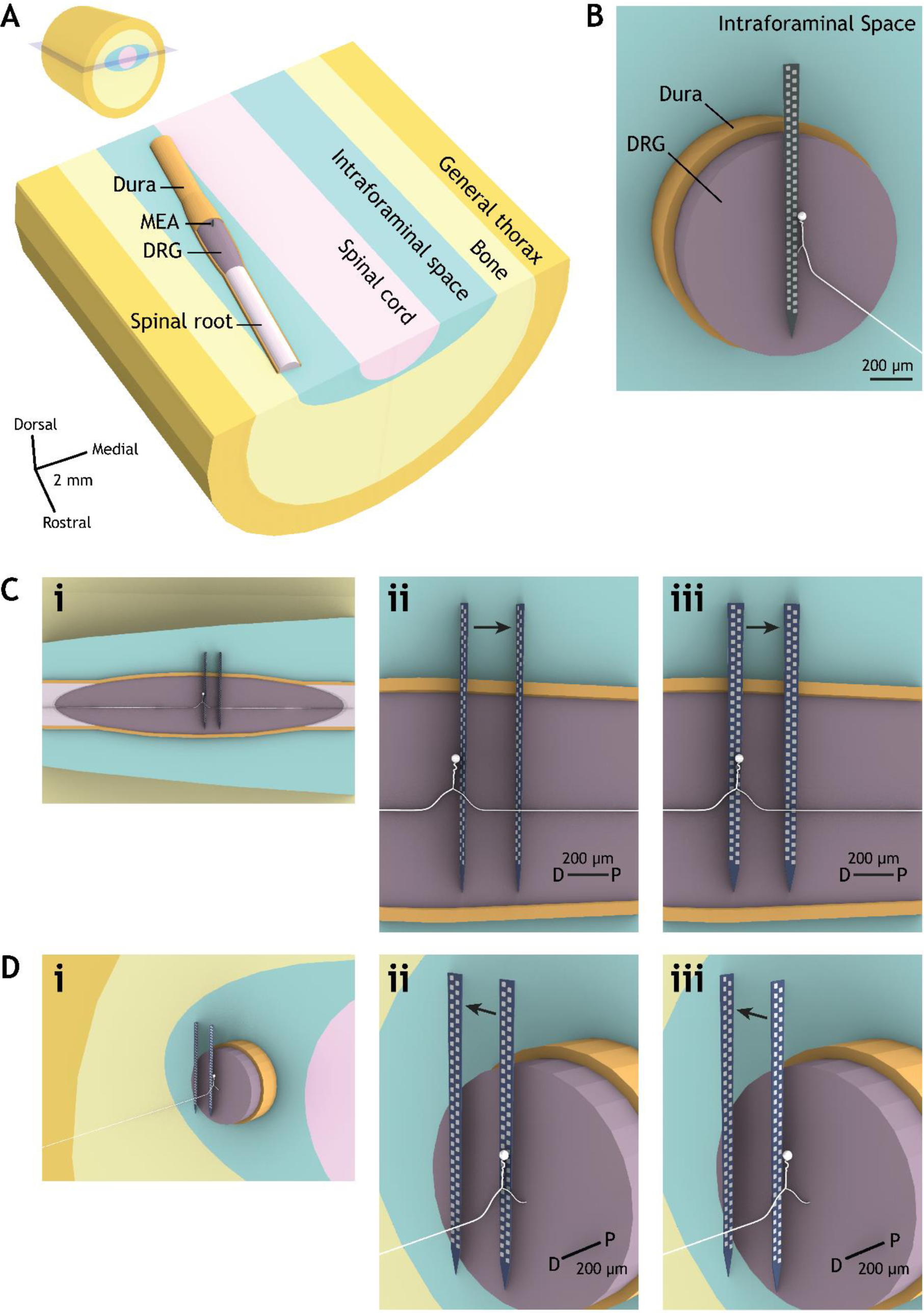
FEM model geometry. (A) The full FEM model. The light blue surface in the inset in the upper left is a coronal plane with which the larger, labeled figure is transected. The axis bars are each 2 mm. MEA = microelectrode array. (B) Transverse cross section of the DRG, exposing the MEA. The relative location of the NEURON model is shown in white. (C) Two sets of simulations varied the position of the array along the central branch. The nearest and furthest position of the array from the soma of the NEURON model for these simulation sets are each shown in (ii) and (iii). The array positions where the contacts faced distally along the branch axis are shown in (ii), and array positions where the contacts faced the DRG neuron are shown in (iii). The relative positioning within a longitudinal cross section of the DRG is shown in (i). D = Distal, P = Proximal. (D) The same as in C, but for two sets of simulations where the position of the array varied laterally in the direction perpendicular to the NEURON model’s branches. Array positions where the contacts faced distally along the branch axis are shown in (ii), and array positions where the contacts faced the soma are shown in (iii). The relative positioning within a transverse cross section of the DRG is shown in (i).

**Table 1:** FEM model dimensions.

| Parameter | Value | Source |
| --- | --- | --- |
| DRG length | 4 mm | * |
| DRG width | 1.1 mm | * |
| Sacral proximal root length | 3.2 mm | * |
| Sacral proximal root radius | 0.4 mm | * |
| Sacral angle to midline | 9.2° | * |
| Sacral spinal cord diameter | 3.4 mm | * |
| Dural sheath thickness | 53 μm | [32] |
| Vertebrae pedicle thickness | 2.59 mm | [33] |
| Vertebrae centrum height | 6.5 mm | [33] |
| Vertebrae width | 12.78 mm | [33] |
| Sacral foramina width | 8.66 mm | [33] |
| Sacral foramina height | 4.01 mm | [33] |

\*Collected from unpublished data from prior *in vivo* experiments

**Table 2:** FEM model conductivities.

| Parameter | Value used | Source |
| --- | --- | --- |
| Grey matter (standard value) | 0.26 S/m | Adapted from [34]* |
| White matter | 0.57 S/m (longitudinal),<br>0.083 S/m (transverse) | [35] |
| DRG and root sheath | 0.68 S/m | [36] |
| Bone | 0.02 S/m | [34], [37] |
| General tissue | 0.23 S/m | Average of mammalian thorax values in [34] |
| Fat (intraforaminal space) | 0.038 S/m | Average of mammalian fat values in [34] |
| Polyimide lead body | 6.66e-16 S/m | [38] |
| Platinum electrode contact | 8.9e6 S/m | [28] |

\* Multiplied cat cortex value by ratio of rabbit grey matter to rabbit cortex

DRG are heterogeneous in their conductive components, containing neural cell bodies, axons, and glial cells. For sets of simulations varying the trajectory of the glomerulus and implementing auto-ephaptic coupling (as described in the following section *Neuron Models*), we varied the conductivity of the grey matter DRG. We used three values for feline neural tissue (*Table 2*): 0.083 S/m (the transverse conductivity of white matter), 0.26 S/m (the conductivity of grey matter), and 0.57 S/m (the longitudinal conductivity of white matter). The transverse and longitudinal white matter conductivities served as physiologically relevant minimum and maximum conductivities for DRG grey matter.

For simulations in which we varied the location of the microelectrode array within the DRG, we set the grey matter DRG conductivity to a standard value of 0.26 S/m. In these simulations, we shifted the location of the array within the DRG in four sets of simulations (see *Figure 2C* and *2D*). For the first two sets, we shifted the array to six locations that varied proximally towards the spinal cord, in the direction parallel to the central branch and away from the soma of the neuron model (*Figure 2C*). For the other two sets, we shifted the array away from the soma to six locations that varied laterally from the axis of the DRG, perpendicular to the axis of the peripheral and central branches (*Figure 2D*). Additionally, one of each of these sets had the electrode contacts facing distally along the axis of the neuron’s branches (*Figure 2Cii* and *Figure 2Dii*). The other of each set had the contacts in the perpendicular direction, facing the axis of the branches (*Figure 2Ciii* and *Figure 2Diii*). In each of these four simulation sets, the array started 15 µm from the glomerulus of the neuron model and was shifted to 25, 50, 100, 200, and 300 µm in each respective direction.

For all FEM simulation sets across different glomerulus trajectories, grey matter conductivities, and array positions, we simulated each of six adjacent electrode contacts within one column of the array as the active recording contact. The six contacts together spanned the base of the stem axon to the soma. The different contacts captured the extracellular voltage waveform as it would be recorded near different components of the neuron. For each individual FEM simulation, we set all inactive contacts as equipotential with no net current across their surface, and we set the outer boundaries of the general thorax domain to ground. We applied a load boundary condition of 1 A to the active contact and we solved the electrostatic model in COMSOL using the conjugate gradient method to solve Laplace’s equation:

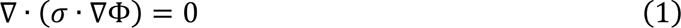

where *σ* is a matrix of the different tissue conductivities and Φ is the resulting potential field.

### Neuron Models

We constructed and simulated the multi-compartment Aδ-LTMR model, and its geometric iterations, in NEURON based on the Aδ-LTMR model described by Graham et al. 2021 [29], [39]. To create a prototypical, anatomically detailed glomerulus of an Aδ neuron, we traced the trajectory of a stem axon and glomerular initial segment (GIS) in Rhino® (Robert McNeel & Associates, USA) from a drawing of a feline A-type trigeminal pseudounipolar neuron by Ramón y Cajal, 1911 [16], [40]. Extending from the base of the stem axon, we built the peripheral and central axon branch trajectories in Rhino 3D to project laterally and medially, respectively, along the axis of the dorsal spinal roots. We used double cable properties describing morphological and electrical myelination parameters, including those dependent on fiber diameter, from McIntyre et al. 2002 (i.e., the MRG model) [39], [41]. Consistent with feline Aδ fiber dimensions found in the literature, we modeled the peripheral and stem axons as 5.7-µm diameter fibers and the central axon as a 3.0-µm diameter fiber [42]. We modeled the 263-µm long GIS as a 1.2-µm diameter unmyelinated axon. This diameter value was calculated by scaling the 1.9-µm node diameter for 5.7-µm fibers from McIntyre et al. 2002 by the 5-µm to 8-µm ratio of initial segment and node diameter values found in Amir and Devor, 2003 [41], [43]. We represented each myelin sheath in the neuron model as six internodal compartments between two main paranode compartments, which were between two myelin attachment paranode compartments [41]. Stem, peripheral, and central axon myelination is shorter and thinner near the T-junction of A-type pseudounipolar neurons [43], [44]. In our model, we determined variations in myelination lengths and thicknesses near the T-junction by scaling the MRG-based myelination parameters (length and number of lamellae for both internodal and paranodal compartments) with ratios described in Amir and Devor, 2003 [43]. We scaled the lengths for the stem axon myelin with these ratios to fit the length of the stem axon traced from Ramón y Cajal, 1911 [16]. We modeled the soma as a series of disc-shaped compartments stacked in the same direction as the GIS compartment most proximal to the soma. The discs were 1-µm long compartments that varied in diameter to approximate a 40-µm diameter sphere (*Figure 3A*).

**Figure 3:**
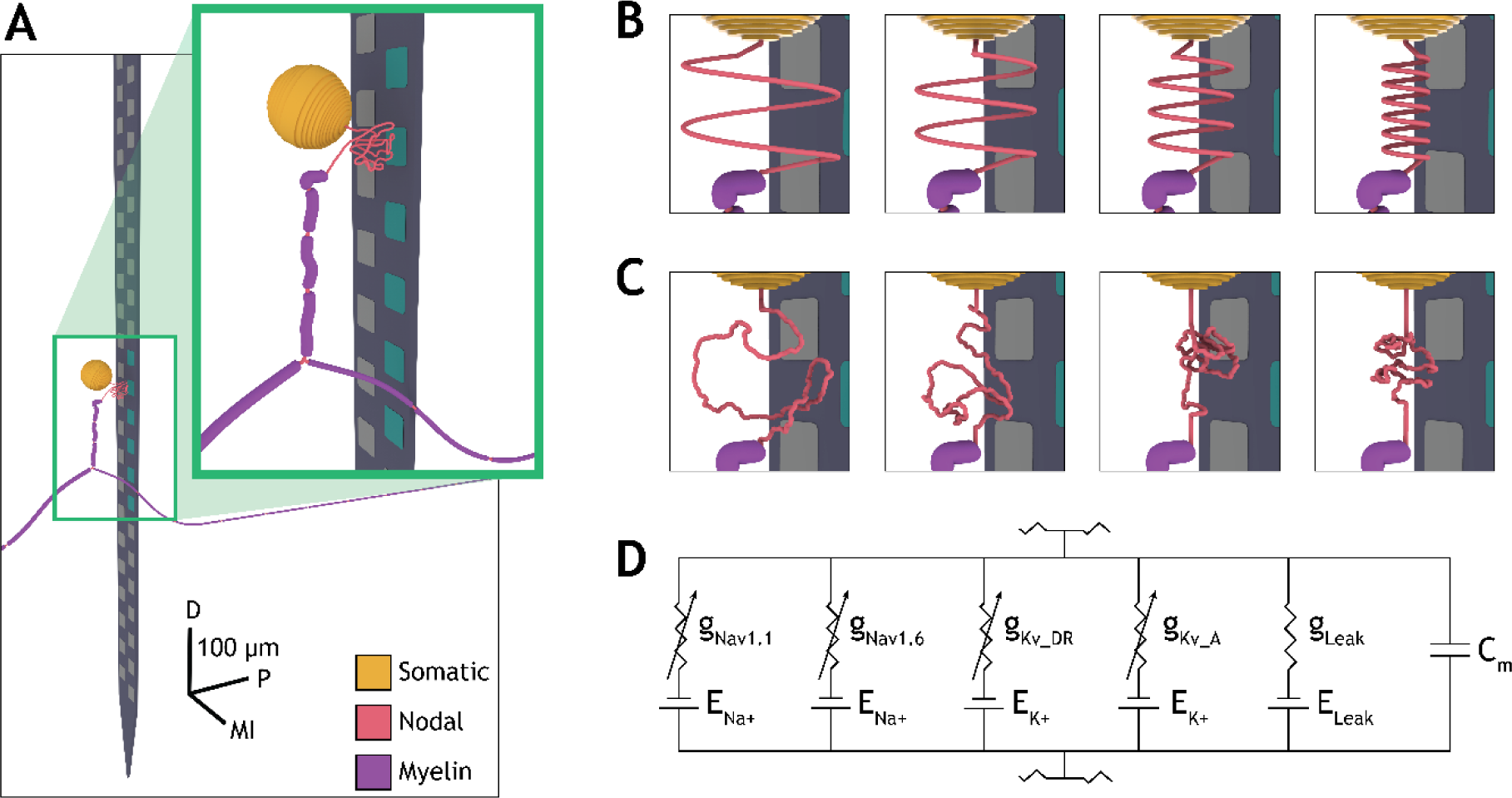
The NEURON model and its variations. (A) A rendering of the NEURON model next to the microelectrode array model. The colored key on the bottom right indicate compartment types. The axis directions are D = Dorsal, P = Proximal, MI = Medioinferior. Teal contacts indicate those set as active recording contacts in the model. (B) Varying spiral trajectories, increasing in tortuosity level (left to right). (C) Varying random walk trajectories, increasing in tortuosity level (left to right). (D) A schematic of the circuit model used in node, GIS, and somatic compartments. See the Neuron Models section and Table 3 for details.

**Table 3:** NEURON model ion channel dynamics.

| **Ion channel** | **Dynamics parameters reference** | **Expression in Aδ-LTMRs reference(s)** | **Maximum conductance ($\bar{g}$)** |
| --- | --- | --- | --- |
| Nav1.1 (TTX-S, transient) | [45] | [45], [55] | Nodes: 0.008 S/cm<sup>2</sup> [45]<br>Soma: 0.0088 S/cm<sup>2</sup> (MCMC)<br>GIS: 0.0088 S/cm<sup>2</sup> (MCMC) |
| Nav1.6 (TTX-S, transient) | [46] | [45], [55], [56] | Nodes: 3.0 S/cm<sup>2</sup> [46]<br>Soma: 0.23 S/cm<sup>2</sup> (MCMC)<br>GIS: 0.3 S/cm<sup>2</sup> (MCMC) |
| K<sub>DR</sub> (Delayed rectifier, slow inactivating) | [47] | [45], [57] | 0.0056 S/cm<sup>2</sup> (MCMC) |
| K<sub>A</sub> (A-type, fast inactivating) | [47] | [45], [58], [59] | 0.112 S/cm<sup>2</sup> (MCMC) |
| Leak | n/a | n/a | Axons: 0.006 S/cm<sup>2</sup> [41]<br>Soma: 0.0018 S/cm<sup>2</sup> (MCMC)<br>GIS: 0.0019 S/cm<sup>2</sup> (MCMC) |

TTX-S: tetrodotoxin sensitive

**Table 4:** NEURON model validation metrics.

| **Parameter** | **Literature Value** | **Source** | **Model Value** |
| --- | --- | --- | --- |
| Resting membrane potential | -51.1 ± 6.1 mV | [42] | -48.9 mV |
| Soma action potential amplitude | 75 ± 8.7 mV | [60] | 77.0 mV |
| Action potential duration (resting to base) | 1.76 ± 0.28 ms | [61] | 1.55 ms |
| Rise time | 0.69 ± 0.094 ms | [61] | 0.69 ms |
| Fall time | 1.07 ± 0.22 ms | [61] | 0.85 ms |
| Afterhyperpolarization duration to 80% | 5.7 ± 6.6 ms | [62] | 6.0 ms |
| Peripheral branch conduction velocity | 2-36 m/s | [60] | 11 m/s |
| Central branch conduction velocity | 1.3-12 m/s | [62] | 4 m/s |

We generated additional simulated GIS trajectories that were each 200-µm long to characterize how the tortuosity of the GIS influenced the recorded extracellular voltage waveform. One set of simulated trajectories implemented nine different spiral paths, where we increased the tortuosity across the set of paths by decreasing the width of the spirals and increasing the number of spiral turns for each trajectory. The spiral paths had identical lengths and terminated at the same global coordinates, so we could construct the soma in the same position relative to the microelectrode array across all trajectories (*Figure 3B*).

In a second set of simulated trajectories of identical length, we implemented a novel random-walk algorithm. A full description of this algorithm is available in the Supplemental Methods section, which includes an online source for our Python code that generates the trajectories. This algorithm ensured that all random walk trajectories were constructed with the same path length and also that the soma was built in the same position relative to the stem axon and the microelectrode array. Example generated trajectories are shown in *Figure 3C* and *Figure S3*.

Lastly, for further comparison, we built a straight GIS path of 32 µm in length. While this pathway was shorter, and thus contained fewer compartments than those of the 200-µm long spiral and random walk trajectories, we maintained the location of the soma relative to the recording contacts. Further, we also ran a set of simulations in which we only varied the length of the GIS. We built spiral trajectories with the same spiral width but different numbers of spiral turns for GIS lengths of 50, 100, 150, 200, and 250 µm.

The nodal, GIS, and somatic compartments included Nav1.1 and Nav1.6 sodium channel dynamics in addition to delayed rectifier and A-type potassium channel dynamics [45]–[47]. Peripheral, central, and stem axon nodes had a Nav1.1 maximum conductance (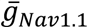) of 0.008 S/cm^2^ [45], a Nav1.6 maximum conductance (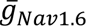) of 3.0 S/cm^2^ [46], and a passive leak conductance (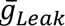) of 0.006 S/cm^2^ [48]. We estimated the potassium channel maximum conductance values (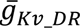 and 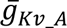) for all compartments with active membrane properties, in addition to sodium and leak conductance values for the soma and GIS, using the Affine-Invariant Markov Chain Monte Carlo (MCMC) method via the *emcee* Python package [49], [50] (*Table 3*). First implemented for this purpose by Adams et al. 2018 [51], we used this method to identify maximum conductance values that produced a somatic intracellular action potential voltage profile (ICAPVP) resembling that of D-hair type Aδ-LTMR neurons found in the literature [51], [52]. This application of the MCMC method uses Bayes’ theorem to estimate the probability distributions for each given parameter and covariances between the parameters to optimize their values in reference to experimental data. We set *a priori* boundaries for conductance values that were generated and tested by the MCMC algorithm. Conductances had to be greater than zero. They also had to be within physiologically relevant ranges (e.g., potassium channel conductances had to be lower than sodium channel conductances). We set the Nav1.6 maximum conductance values in the somatic and GIS compartments used to initialize the MCMC algorithm to 0.15 S/cm^2^ and 0.3 S/cm^2^, respectively. We chose these relative conductances because somata tend to have much lower sodium channel densities than axonal nodes [53], and the soma and GIS generated spontaneous action potentials when the sodium maximum conductance (a proxy for channel density) was too high in these regions [54]. Given an initial set of conductance values and literature-based somatic ICAPVP metrics, we used the MCMC algorithm to test over 150,000 iterations of the model using different values within set ranges of the parameter space. We used several different parameter space sizes – as small as within 10% of the initialization values, and as large as within 300% of the initialization values. For all compartments, we obtained reversal potential values for active membrane dynamics (*E_ion_*) from the sources listed in *Table 3*, and we set the reversal potential value for the passive leak channel (*E_Leak_*) to −52 mV. The resulting maximum conductance parameters that we used in the model are listed in *Table 3*.

### NEURON Simulations

We computed 30 ms of simulated time in NEURON using timesteps of 0.005 or 0.002 ms (for simulations that did or did not incorporate auto-ephaptic coupling, respectively). During each simulation, we used the NEURON `IClamp` function to induce an action potential in the Aδ-LTMR model by applying a 1 nA, 0.2 ms intracellular current pulse to the distal end of the peripheral axon 10 ms after simulation onset.

### Cylindrical Source Summation Method

For NEURON simulations employing auto-ephaptic coupling within the GIS, we used a cylinder source approximation method for calculating extracellular potentials and a mean-field method for calculating the respective ephaptic currents [63]. Each compartment of the neuron was treated as a cylindrical current source, wherein the extracellular potential around a compartment was averaged around the boundary of the compartment. This method makes two assumptions. First, the extracellular space is infinite. Derivations of the point current source equation, such as the cylinder source approximation equation we used here [63], [64]:

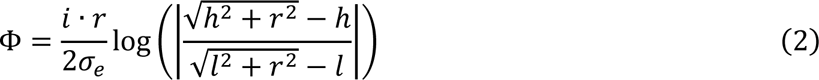

share this assumption. Here, Φ is the potential generated at a field point by a cylindrical compartment, *i* is the current generated by the compartment, *h* and *l* are distances between the field point and either end of the cylindrical source, *r* is the radius of the compartment, and *σ_e_* is the extracellular conductivity around the compartment (equal to the relevant grey matter conductivity value). The domain of ephaptic interactions we simulated was significantly smaller than the extracellular tissue volume modeled around the neuron, which allowed this first assumption to hold. The second assumption is that the extracellular potential at a compartment can be averaged over the boundary of the compartment. Anastassiou et al. 2010 found that in inhomogeneous electric fields, the characteristic length of the field must be at least comparable with or smaller than the cable length in order for the extracellular potential to have a strong effect on the membrane potential [65]. Further, averaging the extracellular potential over the boundary of a compartment requires that the radius of a compartment (*a*) is small relative to the distance between the center axis of the given compartment and the current sources acting on it (*h*), where *a*/*h* << 1 [66]. We built each GIS compartment as 1 micron in length and 1.2 microns in diameter to aim to comply with this restriction, as curves in the GIS trajectory had non-adjacent compartments on the order of only a few microns away from each other.

During each NEURON simulation, after each timestep, we used the membrane current from each compartment along with *Equation 2* to determine the current generated by each compartment and the corresponding extracellular potential generated at all other compartments. We stored these values in a *N x N* matrix, where *N* is the number of compartments throughout the GIS, and we ignored the diagonal of the matrix in the total calculation of auto-ephaptic current (to avoid treating a compartment as a current source on itself). We set each compartment’s extracellular potential to the total of the summed potentials from all other compartments with the `extracellular` method in NEURON. We then included these extracellular potentials in the calculation of the transmembrane currents in the next timestep. Additionally, we used the Crank-Nicolson method, an implicit numerical method, to solve the simulation with timesteps of 0.002 ms (instead of 0.005 ms), which was small enough to produce stable solutions for our primary simulations.

### Computing Extracellularly Recorded Waveforms

After the NEURON simulation was complete, we used a reciprocity-based approach to calculate the time-dependent voltage generated at the recording electrode [23]. As described above, we applied a unit current source (i.e., 1 A) at the individual recording electrode, set all inactive contacts as equipotential with no net current across their surface, grounded the outer model FEM model boundaries (i.e., 0 V), and solved the Laplace equation to obtain the resulting model tissue potentials. We then interpolated the resulting potentials onto each compartment of the Aδ-LTMR model, interpreting this potential as the potential impressed onto the recording electrode by a unit (i.e., 1 A) current source placed at the spatial location of each compartment. We calculated the overall microelectrode recording by superimposing the potentials generated at the recording electrode from the scaled transmembrane currents of each independent compartment. We performed this process for each of the simulation conditions. Finally, we downsampled the resulting waveform from its simulated frequency (200,000 or 500,000 Hz) to 30,000 Hz and then we applied a high-pass filter at 250 Hz using the SciPy *signal* Python package [67] to simulate experimental recording conditions [14], [68].

## Results

### Electrophysiological Properties

In order to produce an electrophysiologically detailed Aδ neuron model, we sought to (1) determine ion channel maximum conductance values that best replicated somatic ICAPVPs found in the literature and (2) determine if accounting for auto-ephaptic coupling within the glomerulus of A-type pseudounipolar neurons produces more physiologically realistic results for action potential propagation behaviors.

We implemented an MCMC algorithm with a model that included the prototypic version of the GIS, traced from Ramón y Cajal 1911 [16], to estimate ion channel maximum conductance values that we would use for subsequent NEURON simulations. The values used in the model, which can be found in *Table 3*, were the most optimal values produced by the algorithm and produced somatic ICAPVPs that sufficiently matched both quantitative metrics and qualitative properties of voltage profile shapes found in the literature. An example somatic action potential is shown in *Figure 4A*. Contour plots generated from the MCMC optimization can be found in *Figure S1*.

**Figure 4:**
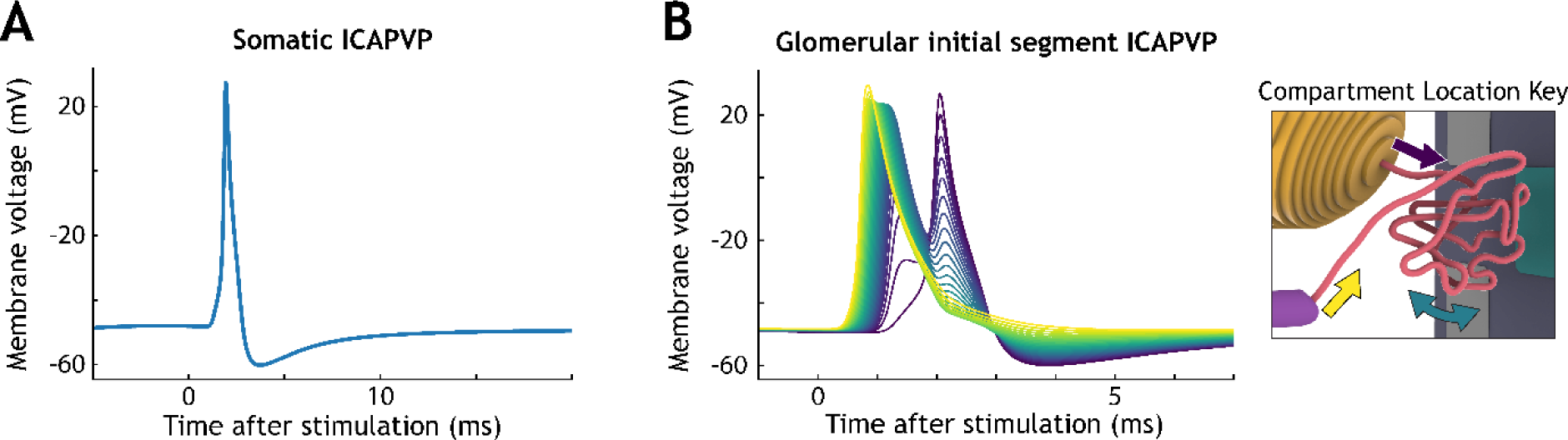
Intracellular action potential voltage profiles (ICAPVPs) of the soma and GIS. (A) The ICAPVP of the center somatic compartment. (B) The ICAPVP of GIS compartments at intervals of ten compartments across the 263 compartments in the GIS. The colors of the lines correspond with the location of the compartment, as indicated by the key on the right.

Using the prototypic version of the NEURON model, we ran an initial simulation without implementing auto-ephaptic coupling to observe the intracellular voltage behaviors. The GIS ICAPVP under traditional simulation conditions is shown in *Figure 4B*. Within the GIS compartments, a back-propagation effect is seen, where the action potential rebounds from the soma back towards the T-junction. This M-shaped ICAPVP is also seen in the T-junction of other DRG neuron models [69], [70]. Next, we ran simulations with varying extracellular conductivities (i.e., DRG gray matter conductivity) while implementing auto-ephaptic coupling. These simulations produced ICAPVPs that were virtually identical to those that did not include auto-ephaptic coupling (*Figure S2A and B*). We ran additional simulations using sweeps of extracellular conductivity values to try and identify a critical range in which auto-ephaptic interactions might notably perturb the neuron model’s ICAPVP. However, we found no such range using this method; rather, we identified a cutoff extracellular conductivity value for stability. Simulations using an extracellular conductivity at and below 0.0224 S/m quickly became unstable. This instability was likely due to the timestep resolution that allowed for the completion of simulations within a reasonable timeframe. Alternatively, simulations using an extracellular conductivity as low as 0.0225 S/m did not produce an ICAPVP that was consequentially different from those that did not implement auto-ephaptic coupling (*Figure S2C and D*). Note that we calculated all simulated waveforms shown within figures without implementing auto-ephaptic coupling unless otherwise indicated.

### Neuronal Structure

Next, to observe how the tortuosity of the GIS affects recorded extracellular voltage waveforms, we varied the shape and trajectory of the GIS (*Figure 3B-C*; *Figure S3*). We used a series of spiral-shaped trajectories for the GIS pathway during simulations where the soma was positioned approximately 30 µm from the electrode array. The resulting simulated waveforms are shown in *Figure 5*. Additionally, we used a number of trajectories built with a random walk algorithm to simulate the meandering GIS pathways of many DRG A-type neurons (*Figure S3*). We made thirty iterations of different trajectories for each tortuosity level. While the random trajectories within each tortuosity level were unique, including as compared to the spiral trajectories of the same relative tortuosity, the general qualitative shape of their recorded waveforms were similar in shape to those of the same degree of tortuosity. Waveforms produced by a series of example random walk trajectories are shown in *Figure 6*.

**Figure 5:**
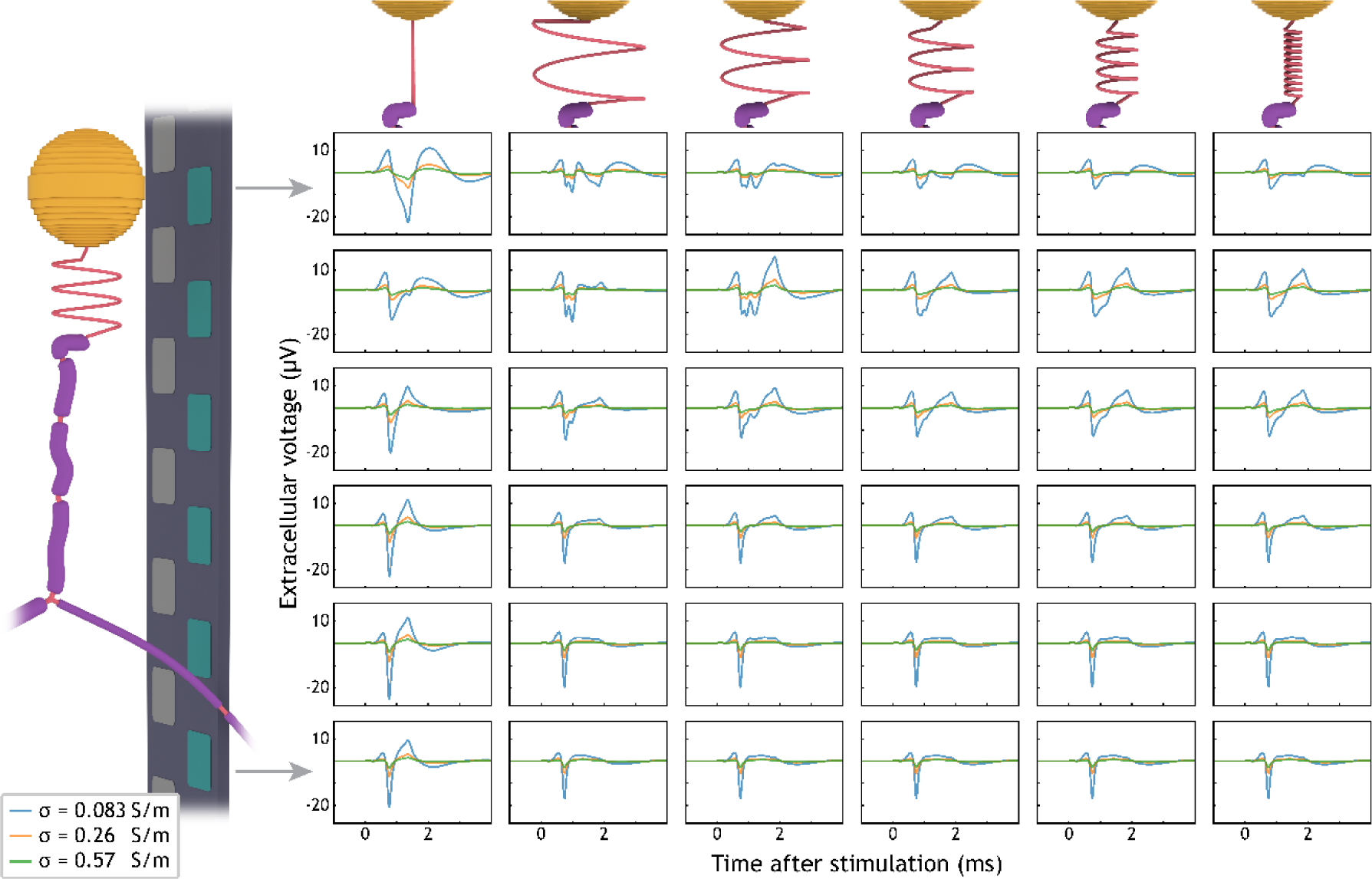
Waveforms recorded from NEURON model variations with spiral GIS trajectories. Each individual plot shows waveforms simulated with different grey matter conductivity values, as indicated by the key on the bottom left. Each row of plots shows waveforms recorded from each electrode contact, as indicated by the schematic on the left. Each column shows waveforms recorded from versions of the NEURON model with the GIS trajectory shown via the rendering at the top of each column. The far-left column shows the 32 µm straight pathway. The subsequent columns show tortuosity levels 1, 3, 5, 7, and 9.

**Figure 6:**
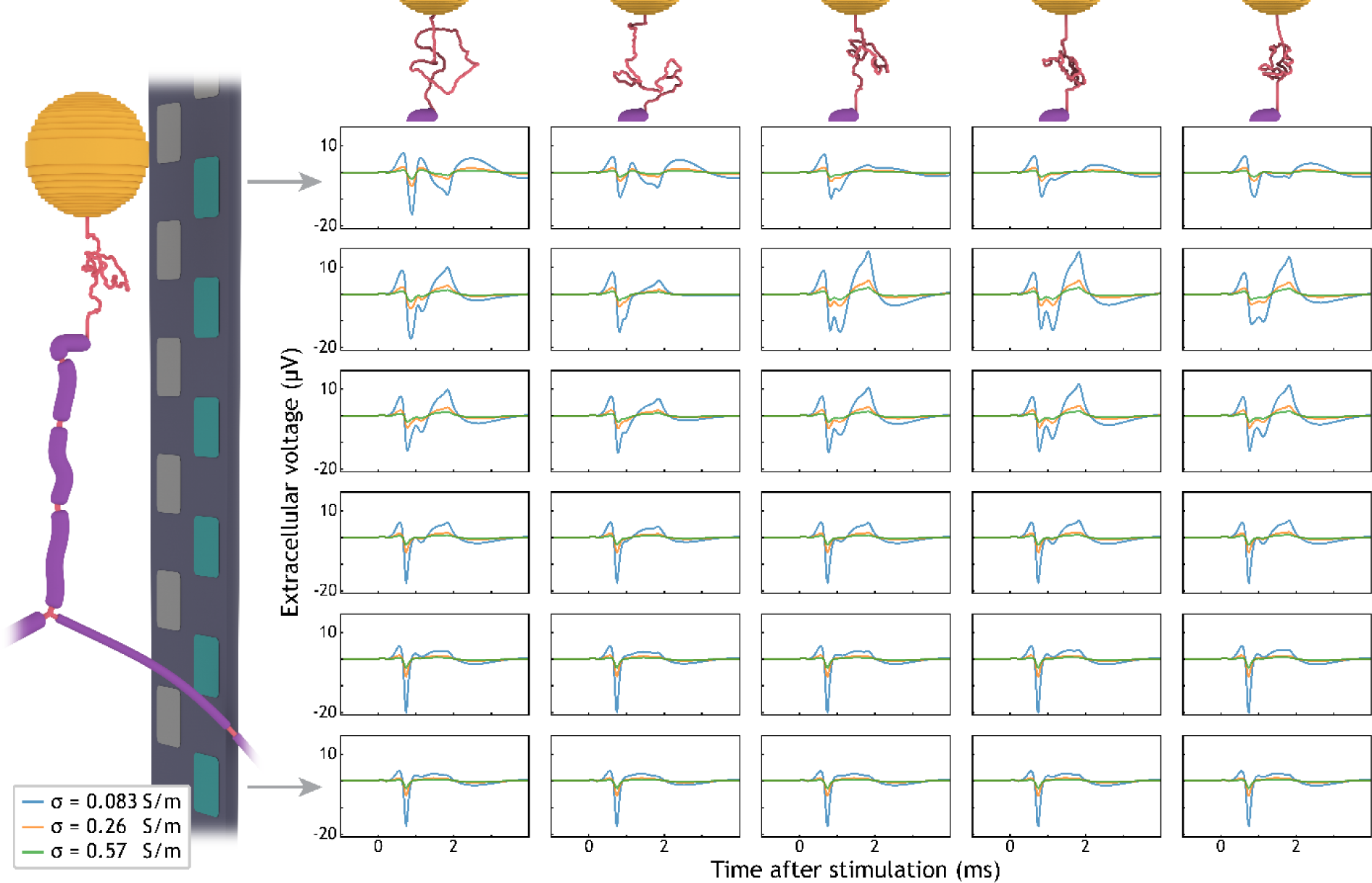
Waveforms recorded from NEURON model variations with example random walk GIS trajectories. See caption for Figure 5. The columns here show random walk trajectories of tortuosity levels 1, 3, 5, 7, and 9.

Simulations using both types of GIS trajectories indicate that those with greater tortuosity produced extracellular voltage waveforms that had different multiphasic properties than those that were less tortuous. Further, the location of the active recording electrode relative to components of the neuron had a significant impact on the general shape of the waveform. We used peak-to-peak voltage amplitude, number of inflection changes, and number of zero crossings as metrics to capture the multiphasic properties of the waveform shapes. Recordings near the T-junction resembled those typically seen recorded near straight axons. Meanwhile, waveform shapes near the soma and GIS significantly differed from each other and recordings near the T-Junction. Waveforms recorded near the soma generally had larger peak-to-peak amplitudes, more inflection changes, and more zero crossings than those recorded near the GIS and T-junction (*Figure 7 and Figure S4*). Waveforms that were recorded on electrodes closer to the T-junction were comprised primarily of current emitted from the peripheral and central branches (*Figure S5*). Likewise, waveforms that were recorded on electrodes closer to the soma and GIS were comprised primarily of current emitted from these components. The large and wide positive voltage deflection seen at the end of waveforms recorded near the soma can be attributed to the large amount of capacitive current that somata emit when they fire (*Figure S5*).

**Figure 7:**
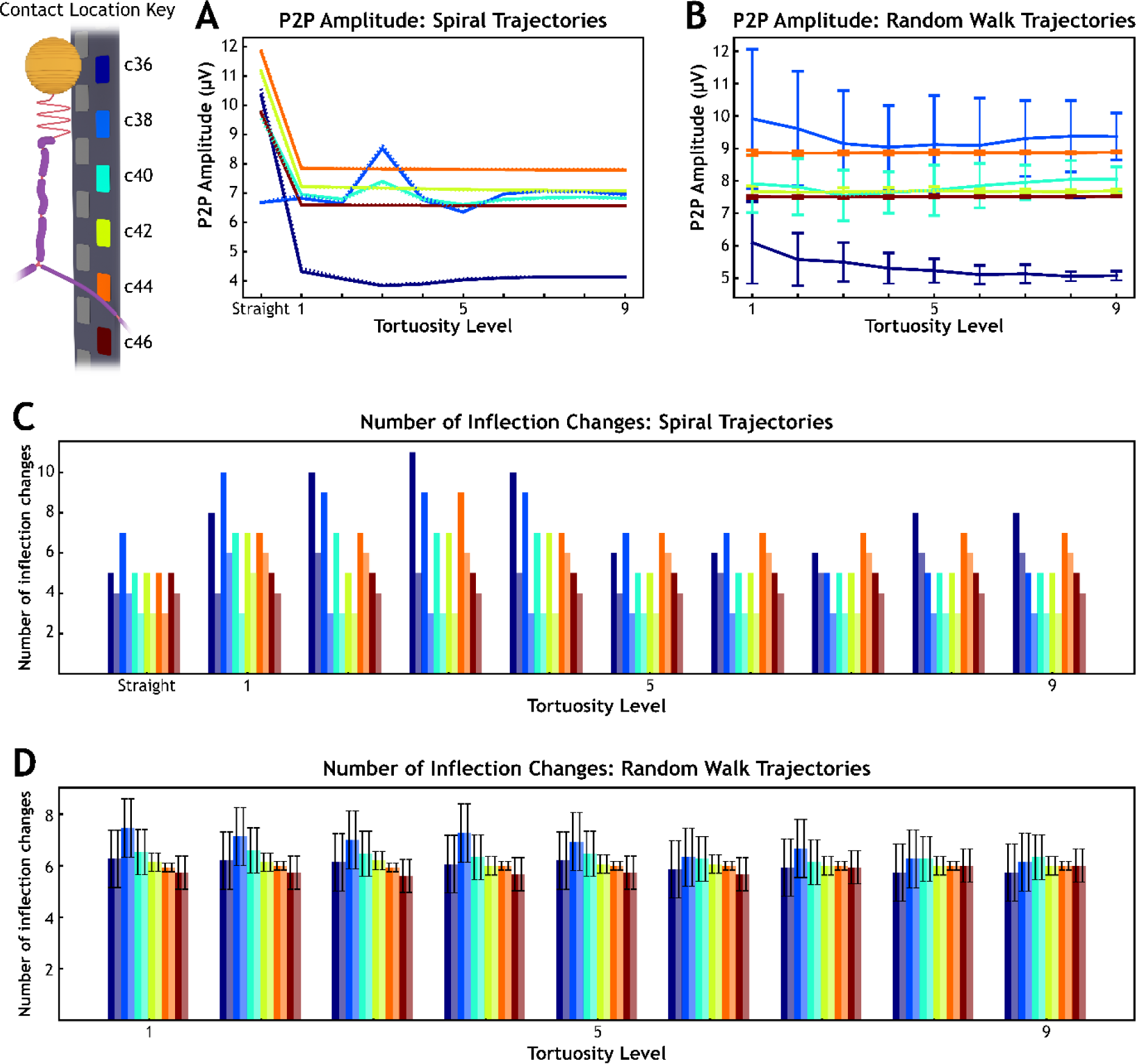
Summary plots of how GIS tortuosity influences recording peak-to-peak (P2P) amplitude and the number of inflection changes in the waveform. A and B) The color of the lines correspond with the color of the contact on the key in the top left. Solid lines represent results for simulations that did not implement auto-ephaptic coupling, and dotted lines (nearly indistinguishable from solid lines) represent results for simulations that did. C and D) The color of the bars correspond with the color of the contact on the key in the top left. Darker and lighter colored bars represent results for simulations without and with auto-ephaptic coupling, respectively.

We ran each of these types of simulations both with and without auto-ephaptic coupling. Most spiral GIS trajectories that we simulated with auto-ephaptic coupling often produced waveforms that had fewer inflection changes and fewer zero crossings than those we did not simulate with auto-ephaptic coupling (*Figure 7C* and *Figure S4A*). However, these differences in multiphasic properties were subtle in the resulting waveforms. Differences between the waveforms that did and did not implement auto-ephaptic coupling were negligible, regardless of GIS tortuosity and extracellular conductivity (*Figure S6*).

As a subsequent attempt at using the cylindrical source summation method, we further divided the GIS compartments into 10 segments each and decreased the timesteps by half to increase the spatial and temporal resolution of the solution. These simulations were run with the lowest physiologically relevant extracellular conductivity value (0.0826 S/m). Further, we tested two GIS trajectories: the most tortuous spiral trajectory and one of the most tortuous random walk trajectories. With these adjustments, the changes in the resulting GIS ICAPVPs were still negligible with respect to those without incorporating auto-ephaptic coupling via the cylindrical source summation method.

Additionally, we performed a sweep of simulations where the spiral width of the GIS trajectory was maintained, and only the length was varied. The waveforms recorded by each contact for each GIS length are shown in *Figure 8*. Our simulations indicated that shorter GIS trajectories resulted in less current being emitted by the GIS, meaning that less current was offsetting the current being emitted by the soma (*Figure S5*). This led to neuron model iterations with shorter GIS trajectories producing extracellular voltages recorded near the soma with greater peak-to-peak amplitudes than iterations with longer GIS trajectories.

**Figure 8:**
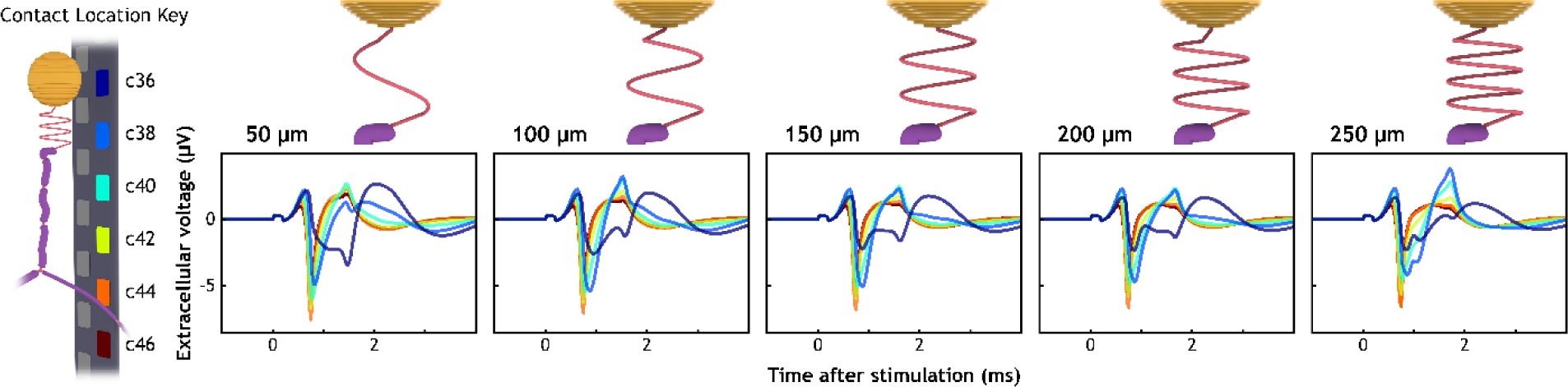
Simulated extracellular recordings of neuron models with different GIS lengths. The different colored lines correspond with different electrode contacts, as indicated by the key on the left. The length of the GIS for each subplot is listed above the plot, next to a rendering of the trajectory.

### Recording Location

Using the prototypic GIS trajectory traced from Ramón y Cajal [16], we simulated recordings of the neuron from two different angles and with the microelectrode array positioned at varying distances in two different directions. *Figure 2C* and *2D* show the angle and positions of the array for these sets of simulations, where the contact face of the array is facing the soma in all positions.

For the two simulation sets where the microelectrode was shifted proximally towards the spinal cord along the central axon, a similar pattern is seen between the simulation with the contacts facing distally along the axis of the branches (*Figure 2Cii, Figure 9A*) and with the contacts facing perpendicular to it (*Figure 2Ciii, Figure 9B*). In both simulation sets, the distance between the recording contacts and the central branch was maintained. Consequently, waveforms that were recorded by contacts nearest the central branch maintained both their general amplitude and shape across simulations for different microelectrode shift distances in this direction. This trend was due to more consistent current contributions of the peripheral and central branches (*Figure S7A and S7B*).

**Figure 9:**
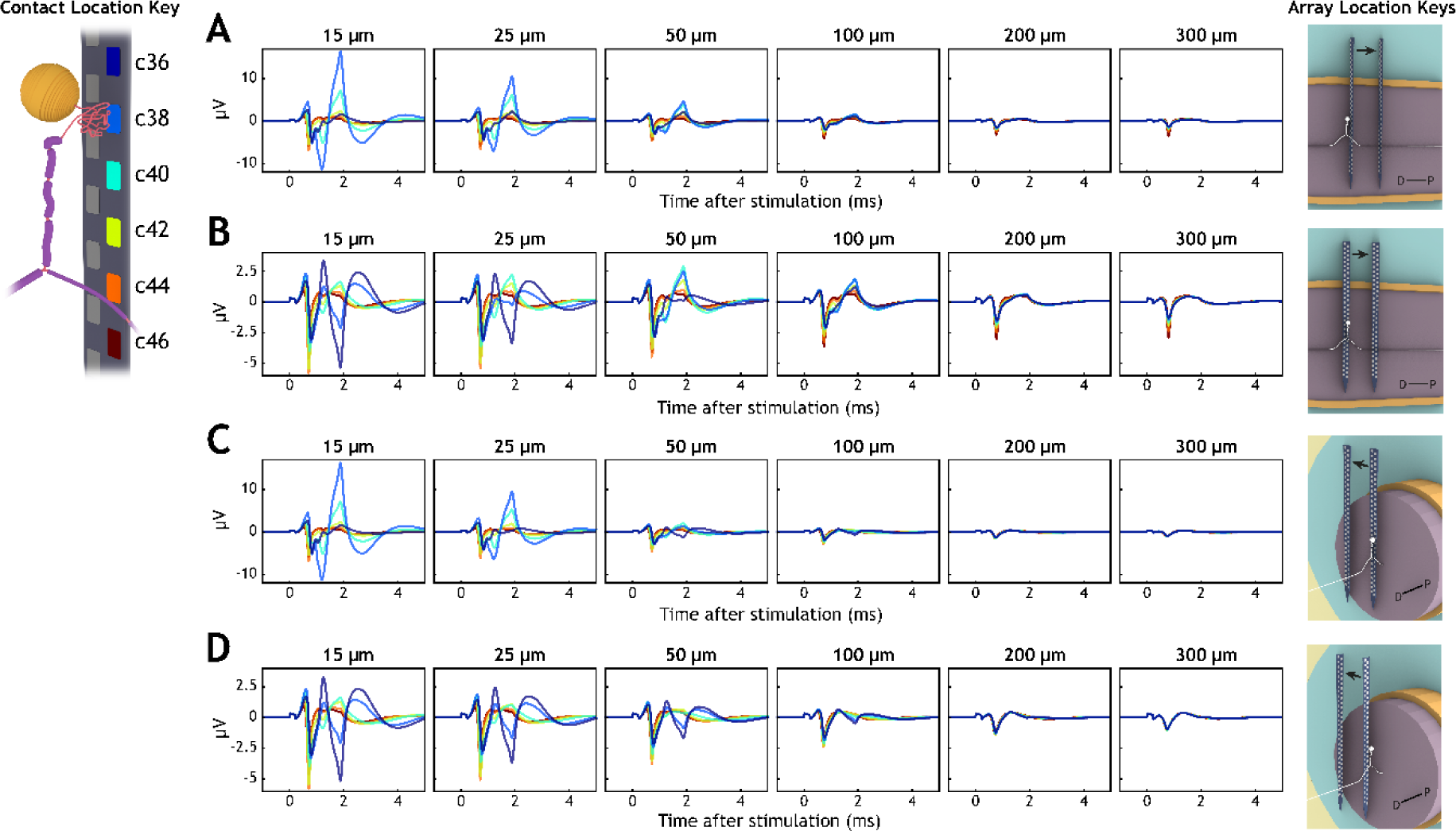
Recorded waveforms from different angles and locations of the array. Waveform colors correspond with the color of the contacts in the key on the left. Distances in each plot subtitle correspond with array positions shown in the key on the right, where the arrow points from the 15-µm position to the 300-µm position. D = Distal, P = Proximal. (A) Corresponds with Figure 2Cii. (B) Corresponds with Figure 2Ciii. (C) Corresponds with Figure 2Dii. (D) Corresponds with Figure 2Diii.

Additionally, simulations that shifted the microelectrode laterally away from the axis of the neuron branches decreased in amplitude. This was true for both simulation sets, with the electrodes facing in both directions, (*Figure 2Dii* shown in *Figure 9C* and *Figure 2Diii* shown in *Figure 9D*). Interestingly, changes in the general shape of waveforms recorded by contacts closer to the GIS and soma (contacts 36 and 38) across different array positions were more prominent in the simulation set where the contacts were facing the axis of the branches (shown in *Figure 9B* and *9D*, as opposed to simulations where the contacts were facing distally along the axis of the branches, shown in *Figure 9A* and *9C*).

## Discussion

Computational modeling of microelectrode recordings has provided a robust understanding of the factors that contribute to the extracellular voltage waveforms produced by neurons in the brain [23]– [25]. In this study, we sought to elucidate the biophysical sources of the distinct extracellular voltage waveform shapes produced by DRG neurons. We coupled an FEM model and multi-compartment double-cable models to simulate a penetrating, multi-site, recording microelectrode array and a physiologically and anatomically detailed A-type sensory neuron, respectively. This work was the first computational study exploring extracellular microelectrode recordings of DRG neurons that extensively explored the effects of glomerular structure and tortuosity on the recorded waveform. It was also the first computational study to implement auto-ephaptic coupling. We demonstrated how factors, such as microelectrode location relative to specific components of the neuron and GIS tortuosity and length, can influence both the amplitude and multiphasic properties of the extracellularly recorded waveform.

### MCMC methods and membrane dynamics

The neurophysiological detail of the neuron model simulated for this study, an Aδ-LTMR, was of key precedence when constructing and validating its electrophysiological behaviors. We validated our model on its ability to reproduce intracellularly recorded action potential characteristics because intracellular voltage waveform shape directly impacts extracellular voltage waveform shape [71], [72]. Previous models of A-type sensory neurons [39], [48] have been constructed for extracellular stimulation studies, for which similar validation metrics demonstrated the models’ abilities to capture electrophysiological behaviors suitably for exploring the effects of neuromodulation. However, for our recording model, we sought to simulate an ICAPVP that qualitatively resembled corresponding *in vivo*/*in vitro* traces found in the literature as closely as possible, particularly for features such as the broad, rounded, afterhyperpolarization phase of the action potential [61], [62], [73]. We believed accounting for this portion of the intracellular voltage waveform would produce extracellular waveforms with more realistic timescales and shapes in reference to *in vivo* recordings.

The use of MCMC methods to estimate ion channel parameter values for a neuron model to optimally reproduce experimental intracellular action potential data was first implemented by Adams et al. 2018 [51]. More recently, Lee et al. 2022 used MCMC methods to create populations of distinct, but biophysically realistic, interneuron models [52]. Here, we used an MCMC algorithm to select conductance values for membrane dynamics that best reproduced the shape of Aδ-LTMR somatic action potentials found in the literature, similar to Adams et al. 2018. In addition to maximum conductances for ion channels, we also included somatic and GIS leak conductance values in the MCMC algorithm. Allowing the somatic and GIS leak conductance parameter space to be explored further with the MCMC algorithm contributed significantly to the qualitative fidelity of the neuron model’s ICAPVP relative to experimental recordings found in the literature [61], [62] (*Figure 4*).

### Neuronal structure and glomerulus trajectory

Glomeruli exist in larger, A-type neurons across species and across different sensory ganglia, and their length and tortuosity increase with axon size and phylogenetic complexity [16], [74]. We sought to characterize how the trajectory of glomeruli impacts the shapes of waveforms recorded by microelectrodes near different components of our pseudounipolar neuron model, which we quantified with peak-to-peak amplitude, the number of inflection changes, and the number of zero crossings. Both the tortuosity of the GIS (independent of length, *Figure 5 and 6*) and the length of the GIS (independent of tortuosity, *Figure 8*) had a significant impact on the multiphasic properties of waveforms recorded closer to the soma and GIS. While the number of inflection points for waveforms recorded near the soma and GIS decreased as GIS tortuosity increased (for both spiral and random walk trajectories, *Figure 7C and D*), the number of zero crossings and peak-to-peak voltage amplitudes were not correlated with GIS tortuosity (*Figure 7A* and *Figure S3*). Across all simulations varying the GIS trajectory, waveforms recorded closer to the T-junction generally maintained their shape, because the potentials being imposed onto these recording contacts originated primarily from the peripheral and central branches. Further, simulated waveforms recorded closer to the T-junction also resembled extracellular action potential waveforms that are often found in peripheral axon recordings [75]–[77] (see simulated waveform plots in the fifth and sixth rows of *Figure 5* and *Figure 6*). Additionally, a number of our simulated waveforms recorded closer to the GIS had striking resemblances to the unique W-shaped waveforms that occasionally are found in *in vivo* DRG microelectrode recordings (*Figure 1B* compared with the top three rows of *Figures 5 and 6*).

For simulation sets that only varied the length of the GIS, waveforms recorded near the soma and GIS had different multiphasic properties across different GIS lengths (*Figure 8*). These differences were presumably due to the degree to which the GIS “absorbed” the rebounded action potential from the soma (long glomeruli had more surface area and thus emitted more current during a somatic action potential) and due to the degree to which it mirrored the negative current phase magnitude of the firing soma. This behavior also led to neurons with shorter glomeruli having slightly larger recorded waveforms near the soma than those with longer ones. Less current emitted immediately by the GIS (and entering the myelinated stem axon instead) meant less current to mirror and offset the large amount of current emitted by the soma. Correspondingly, in comparison to the spiral trajectories, the short, straight GIS trajectory produced much larger peak-to-peak waveforms (*Figure 5*). Further, the multiphasic properties of waveforms recorded at different locations relative to this neuron with a straight GIS – locations near the soma and also near the T-junction – did not change drastically across recording locations.

Across these simulation sets varying the GIS trajectory, the microelectrode array location was fixed in place relative to the neuron model, as an array might be in an experimental setting. One should note, for a given trajectory, the profound differences in recorded waveform shapes depended only on recording contact location relative to different microanatomical components of the same neuron. Neural spike data recorded with microelectrode arrays where contacts are particularly small and close together (as with this array we modeled from Sperry 2021 [14]), or data recorded while electrode drift on the order of several microns is anticipated, should be sorted with caution. Spike sorting techniques, which often rely on principal component analysis methods [78], may classify waveforms from the same neuron (perhaps recorded on different but adjacent contacts) into different units due to the waveforms’ overt differences in shape. Misclassification of waveforms can lead to misrepresentation of spike trains for relevant units. Our findings may provide context for research and development of closed-loop stimulation that uses automated spike sorting for machine learning algorithms and that use microelectrode arrays to record DRG neuron targets.

### Glomerulus electrophysiology and auto-ephaptic coupling

As with many evolutionarily conserved structures in biology, it is reasonable to surmise that glomeruli serve a distinct function for pseudounipolar neurons in sensory ganglia. Devor, 1999 suggested glomeruli may aid the propagation of action potentials across the T-junction from the peripheral axon to the central axon by increasing the resistive properties of the stem axon, and thus minimizing shunting by the soma [79]. This speculation is supported by our simulations, as GIS and somatic ICAPVPs showed action potentials being shunted into the soma before bouncing back towards the T-junction (*Figure 4B*). Shortening the GIS trajectory led to soma-induced perturbations in the peripheral and central ICAPVPs just distally of the T-junction (*Figure S8*). Our simulations suggest that the mere length of the glomerulus helps extend the stem axon and separate the reverberation of the firing soma from the sensory signal propagating towards the central nervous system.

In this study, we also considered ephaptic properties of glomeruli with a focus on the dense coiling of their axonal pathways. Typically, in cable neuron modeling, the extracellular conductivity surrounding a neuron is assumed to be large enough such that the extracellular voltage is negligible with respect to the calculation of the membrane voltage [80]. The incorporation of ephaptic coupling intends to account for small, but arguably non-negligible, action potential-driven changes in the extracellular voltage and their subsequent impact on adjacent excitable membranes’ firing properties [22], [63], [81]. In A-type pseudounipolar neurons, the glomerular initial segments are unmyelinated. They also coil in on themselves in such a way where separate sections of the same fiber can be on the order of a few microns away from each other. Experimental studies have demonstrated that ephaptic interactions between neurons, even those separated by more than a few microns, are not always negligible [20], [21], [65]. It would seem intuitive that ephaptic interactions – or in this case, auto-ephaptic interactions – contribute to the electrophysiological behaviors of these neurons. We hypothesized that the dense coiling of these glomerular trajectories might act to dissipate current and dampen the secondary action potential propagating towards the soma.

We used a cylinder-source summation method to calculate changes in extracellular potential due to membrane currents at the end of each timestep, and those potentials were then used to calculate the ephaptic potential surrounding each compartment during the next time step. Our results showed negligible differences in both ICAPVPs and extracellularly recorded waveforms between simulations that used this method and those that did not (*Figure S2* and *Figure S6*). Lowering the extracellular conductivity below what is traditionally considered physiologically relevant also did not cause notable changes in the simulated intra- and extracellular potentials. At and below a critical extracellular conductivity value, 0.0224 S/m (a value which we found depended on membrane dynamics parameters), the low extracellular conductivity caused the simulation to fail to converge. We tried using smaller time and space discretization, separately and simultaneously. In these cases, either the simulated results did not change, or the simulation was too computationally expensive.

While the methodology we used here to implement auto-ephaptic coupling did not reveal any substantial changes in results compared to traditional simulation methods, using a resistor network method instead may be worth exploring. Resistor networks are capable of providing more computationally rigorous results, but at the cost of being vastly more computationally expensive [81]. The cylinder-source summation method we used here provided computational efficiency and identified parameter ranges that may have biophysical relevance for future studies. Further exploration of auto-ephaptic interactions could have implications for the electrophysiology of other neurons with complex dendritic and axonal geometries, such as cerebellar Purkinge neurons and cortical pyramidal neurons. It could also provide insight for research and computational models of DRG stimulation. Understanding the intricacies of the manner in which DRG neurons are electrically excitable – both with and without the context of extracellular stimulation – will hopefully help further elucidate the underlying mechanisms of neuromodulation therapies that target DRG.

### Recording location and electrode distance

In a subset of simulations in this study, we varied the position of the microelectrode array with respect to the neuron model, shifting it within the mediolateral-rostrocaudal plane while also simulating two directional positions for the contact face of the array (*Figure 2C* and *2D*). Our study showed that when an electrode was positioned farther away from the pseudounipolar neuron, certain features of the waveform – such as the contributions of the peripheral and central branches – were conserved more than others (contact 44 and contact 46 recordings in *Figure 9*). Additionally, we maintained the distance of the array from the main components of the neuron but had the array face in two different directions. Between these sets of simulations, the contact directionality of the array constituted minimal changes for waveform shapes recorded near the branches but significant changes for the multiphasic properties and peak-to-peak amplitude of the waveforms recorded near the soma and GIS (*Figure 9A* versus *9B,* and *Figure 9C* versus *9D*). Other computational studies of single-neuron microelectrode recordings have made similar types of broad conclusions in regard to location-based recording factors [25], [82]. While the direction of the array influenced the overall shape of the waveform, recording distance only influenced the amplitude, and the general shape of the waveform was largely maintained.

Researchers using functional electrophysiology to study the DRG might use this understanding of the distinct types of waveform shapes that are recorded near different components of a pseudounipolar neuron to help with functional mapping. One could record signals from the DRG with microelectrode arrays and distinguish between units whose axonal components, versus the cell body, are near the recording microelectrode. Further, one might identify what proportion of recordings indicate the microelectrode is near a cell body and glomerulus. After all, not all A-type DRG neurons have glomeruli, so future studies might investigate which functionalities are associated with specific geometric phenotypes of neurons.

### Limitations and future directions

One of the most restrictive aspects of creating a functionally detailed Aδ-LTMR neuron is the current paucity of microscale biophysical mapping of these neurons. Experimental compilation of the density and change dynamics of functionally inserted ion channels in different regions of neural membranes would provide quantitative and spatially relevant data on the maximum ion conductivity values for A-type neuron models. Separately, our model simulated waveforms with peak-to-peak voltage amplitudes that were smaller than what is typically seen experimentally. *In vivo*, extracellular DRG recordings are on the order of 30-120 µV, while assuming we are detecting neurons within 200 µm of the electrode contact [72]. The waveforms produced by versions of our Aδ-LTMR model using a grey matter (i.e., extracellular) conductivity value of 0.26 S/m only reached as large as about 30 µV, but they normally were on the order of about 10 µV (within 20 µm between the contact and the GIS, *Figures 5, 6, and 7*). Meanwhile, some somatic current contributions, which were canceled out by nearly identical yet inverse rebounding GIS current contributions, were as large as 50 µV (*Figures S5 and S7*). While these separate voltage amplitude contributions of the soma and GIS are on the order of what is found experimentally for extracellular recordings of these neurons, this effect needs to be explored and validated experimentally. Further, Bestel et al. 2021 found that modeling a dense layer of passive glial cells around the recording target neuron greatly increases the amplitude of extracellular recordings to levels found experimentally [25]. DRG contain numerous satellite glial cells that surround the primary sensory somata [83], [84]. While our model captured the complex qualitative features of DRG neuron recordings, it is reasonable to conclude that incorporating a satellite glial layer would lead to simulated recording amplitudes that more accurately represent what is found *in vivo*.

Future computational studies of sensory neuron modeling and neural recording modeling could take many forms. As we have discussed here, the biophysical properties of sensory neuron models could be represented more accurately in several ways. Further ion channel localization research may lead to more accurate representations of the currents that contribute to action potential propagation throughout different parts of these neurons. A broad analysis of how recorded waveforms of different sensory neuron types, particularly at different distances from the recording electrode, compare to each other would also be beneficial for peripheral electrophysiology research. This analysis may provide insights as to how the shape of an extracellularly recorded waveform changes with respect to neuron type (i.e., axon size and ion channel expression profile) and distance from a recording microelectrode. In turn, this insight might inform neuron type classification based only on extracellularly recorded waveforms.

Lastly, the waveform properties of DRG neuron microelectrode recordings might have embodied different degrees of change in response to the factors explored here if a different microelectrode was modeled. Utah array electrodes can have surface areas that are larger than the contacts on the microelectrode array modeled for this study [14], [85], [86], which could influence both the shape and span of different multiphasic properties of the recorded waveform. We anticipate that factors explored here, such as neuron geometry and electrode location relative to the neuron, would still be relevant.

However, future studies modeling different types of electrode contact shapes and sizes are needed to further explore these effects.

## Conclusions

Our study utilized a computational modeling approach to provide insight and context for electrophysiology studies using microelectrodes to record the activity of pseudounipolar neurons at the DRG. Our results demonstrate that researchers can make reasonable inferences about which general region of a pseudounipolar neuron a recording microelectrode contact is near based on the qualitative shapes of its extracellular waveform. Further, the contact size, count, and pitch of microelectrode arrays should be considered when spike sorting these types of neural recordings, whether it is being performed manually or by a machine-learning-based classification algorithm. Our model illustrated how waveforms from the same neuron can look entirely different depending on which microanatomical components the recording contact is near. As we have demonstrated in this computational study, there are many factors that can influence the unique and diverse shapes of extracellular waveforms recorded from DRG A-type neurons.

## Acknowledgments

This study was funded by the NSF CAREER Award 1653080. NEURON model simulations were executed in part via services from Advanced Research Computing at the University of Michigan, Ann Arbor.

## References

[1] C. T. Nordhausen, E. M. Maynard, and R. A. Normann, “Single unit recording capabilities of a 100 microelectrode array,” Brain Research, vol. 726, no. 1, pp. 129–140, Jul. 1996, doi: 10.1016/0006-8993(96)00321-6.

[2] H. Takahashi, M. Nakao, and K. Kaga, “Cortical mapping of auditory-evoked offset responses in rats,” NeuroReport, vol. 15, no. 10, pp. 1565–1569, Jul. 2004, doi: 10.1097/01.wnr.0000134848.63755.5c.

[3] T. I. Shiramatsu, K. Takahashi, T. Noda, R. Kanzaki, H. Nakahara, and H. Takahashi, “Microelectrode mapping of tonotopic, laminar, and field-specific organization of thalamo-cortical pathway in rat,” Neuroscience, vol. 332, pp. 38–52, Sep. 2016, doi: 10.1016/j.neuroscience.2016.06.024.

[4] S. N. Flesher et al., “A brain-computer interface that evokes tactile sensations improves robotic arm control,” Science, vol. 372, no. 6544, pp. 831–836, May 2021, doi: 10.1126/science.abd0380.

[5] G. T. A. Kovacs et al., “Silicon-substrate microelectrode arrays for parallel recording of neural activity in peripheral and cranial nerves,” IEEE Transactions on Biomedical Engineering, vol. 41, no. 6, pp. 567–577, Jun. 1994, doi: 10.1109/10.293244.

[6] A. Branner and R. A. Normann, “A multielectrode array for intrafascicular recording and stimulation in sciatic nerve of cats,” Brain Research Bulletin, vol. 51, no. 4, pp. 293–306, Mar. 2000, doi: 10.1016/S0361-9230(99)00231-2.

[7] D. J. Weber, R. B. Stein, D. G. Everaert, and A. Prochazka, “Decoding sensory feedback from firing rates of afferent ensembles recorded in cat dorsal root ganglia in normal locomotion,” IEEE Transactions on Neural Systems and Rehabilitation Engineering, vol. 14, no. 2, pp. 240–243, Jun. 2006, doi: 10.1109/TNSRE.2006.875575.

[8] L. E. Tennyson, C. Tai, and C. J. Chermansky, “Using the Native Afferent Nervous System to Sense Bladder Fullness: State of the Art,” Curr Bladder Dysfunct Rep, vol. 11, no. 4, pp. 346–349, Dec. 2016, doi: 10.1007/s11884-016-0391-2.

[9] U. Topalovic et al., “A wearable platform for closed-loop stimulation and recording of single-neuron and local field potential activity in freely moving humans,” Nat Neurosci, vol. 26, no. 3, Art. no. 3, Mar. 2023, doi: 10.1038/s41593-023-01260-4.

[10] Z. Ouyang et al., “Closed-loop sacral neuromodulation for bladder function using dorsal root ganglia sensory feedback in an anesthetized feline model,” Med Biol Eng Comput, vol. 60, no. 5, pp. 1527–1540, May 2022, doi: 10.1007/s11517-022-02554-8.

[11] Y. Aoyagi, R. B. Stein, A. Branner, K. G. Pearson, and R. A. Normann, “Capabilities of a penetrating microelectrode array for recording single units in dorsal root ganglia of the cat,” Journal of Neuroscience Methods, vol. 128, no. 1, pp. 9–20, Sep. 2003, doi: 10.1016/S0165-0270(03)00143-2.

[12] B. J. Holinski, D. G. Everaert, V. K. Mushahwar, and R. B. Stein, “Real-time control of walking using recordings from dorsal root ganglia,” J. Neural Eng., vol. 10, no. 5, p. 056008, Aug. 2013, doi: 10.1088/1741-2560/10/5/056008.

[13] T. M. Bruns, J. B. Wagenaar, M. J. Bauman, R. A. Gaunt, and D. J. Weber, “Real-time control of hind limb functional electrical stimulation using feedback from dorsal root ganglia recordings,” J. Neural Eng., vol. 10, no. 2, p. 026020, Mar. 2013, doi: 10.1088/1741-2560/10/2/026020.

[14] Z. J. Sperry et al., “High-density neural recordings from feline sacral dorsal root ganglia with thin-film array,” J. Neural Eng., vol. 18, no. 4, p. 046005, Mar. 2021, doi: 10.1088/1741-2552/abe398.

[15] S. Matsuda and Y. Uehara, “Cytoarchitecture of the Rat Dorsal Root Ganglia as Revealed by Scanning Electron Microscopy,” J Electron Microsc (Tokyo), vol. 30, no. 2, pp. 136–140, Jan. 1981, doi: 10.1093/oxfordjournals.jmicro.a050295.

[16] S. Ramón y Cajal, Histologie du système nerveux de l’homme & des vertébrés. Maloine, 1909–11, 1911.

[17] R. S. Shivacharan et al., “Neural recruitment by ephaptic coupling in epilepsy,” Epilepsia, vol. 62, no. 7, pp. 1505–1517, 2021, doi: 10.1111/epi.16903.

[18] K.-S. Han, C. H. Chen, M. M. Khan, C. Guo, and W. G. Regehr, “Climbing fiber synapses rapidly and transiently inhibit neighboring Purkinje cells via ephaptic coupling,” Nature Neuroscience, vol. 23, no. 11, Art. no. 11, Nov. 2020, doi: 10.1038/s41593-020-0701-z.

[19] K.-S. Han, C. Guo, C. H. Chen, L. Witter, T. Osorno, and W. G. Regehr, “Ephaptic Coupling Promotes Synchronous Firing of Cerebellar Purkinje Cells,” Neuron, vol. 100, no. 3, pp. 564–578.e3, Nov. 2018, doi: 10.1016/j.neuron.2018.09.018.

[20] R. Amir and M. Devor, “Functional cross-excitation between afferent A- and C-neurons in dorsal root ganglia,” Neuroscience, vol. 95, no. 1, pp. 189–195, Nov. 1999, doi: 10.1016/S0306-4522(99)00388-7.

[21] C.-C. Chiang, R. S. Shivacharan, X. Wei, L. E. Gonzalez-Reyes, and D. M. Durand, “Slow periodic activity in the longitudinal hippocampal slice can self-propagate non-synaptically by a mechanism consistent with ephaptic coupling,” The Journal of Physiology, vol. 597, no. 1, pp. 249–269, 2019, doi: 10.1113/JP276904.

[22] M. Capllonch-Juan and F. Sepulveda, “Modelling the effects of ephaptic coupling on selectivity and response patterns during artificial stimulation of peripheral nerves,” PLoS Comput Biol, vol. 16, no. 6, p. e1007826, Jun. 2020, doi: 10.1371/journal.pcbi.1007826.

[23] M. A. Moffitt and C. C. McIntyre, “Model-based analysis of cortical recording with silicon microelectrodes,” Clinical Neurophysiology, vol. 116, no. 9, pp. 2240–2250, Sep. 2005, doi: 10.1016/j.clinph.2005.05.018.

[24] S. F. Lempka, M. D. Johnson, M. A. Moffitt, K. J. Otto, D. R. Kipke, and C. C. McIntyre, “Theoretical analysis of intracortical microelectrode recordings,” J Neural Eng, vol. 8, no. 4, p. 045006, Aug. 2011, doi: 10.1088/1741-2560/8/4/045006.

[25] R. Bestel, U. van Rienen, C. Thielemann, and R. Appali, “Influence of Neuronal Morphology on the Shape of Extracellular Recordings With Microelectrode Arrays: A Finite Element Analysis,” IEEE Transactions on Biomedical Engineering, vol. 68, no. 4, pp. 1317–1329, Apr. 2021, doi: 10.1109/TBME.2020.3026635.

[26] A. Furniturewalla, P. Rustogi, E. Patrick, and J. W. Judy, “Modeling the Recording of Intraneural Action Potentials with Microelectrodes Using FEM and Point-Source Methods,” in 2019 9th International IEEE/EMBS Conference on Neural Engineering (NER), Mar. 2019, pp. 1191–1194. doi: 10.1109/NER.2019.8717121.

[27] K. Na et al., “Novel diamond shuttle to deliver flexible neural probe with reduced tissue compression,” Microsyst Nanoeng, vol. 6, p. 37, Jun. 2020, doi: 10.1038/s41378-020-0149-z.

[28] “COMSOL Multiphysics®.” COMSOL AB, Stockholm, Sweden. [Online]. Available: www.comsol.com

[29] N. T. Carnevale and M. L. Hines, The NEURON Book. Cambridge University Press, 2006.

[30] D. Daly, W. Rong, R. Chess-Williams, C. Chapple, and D. Grundy, “Bladder afferent sensitivity in wild-type and TRPV1 knockout mice,” J Physiol, vol. 583, no. Pt 2, pp. 663–674, Sep. 2007, doi: 10.1113/jphysiol.2007.139147.

[31] V. P. Zagorodnyuk, M. Costa, and S. J. H. Brookes, “Major classes of sensory neurons to the urinary bladder,” Autonomic Neuroscience, vol. 126–127, pp. 390–397, Jun. 2006, doi: 10.1016/j.autneu.2006.02.007.

[32] A. Kinaci, W. Bergmann, R. LAW Bleys, A. van der Zwan, and T. P. van Doormaal, “Histologic Comparison of the Dura Mater among Species,” Comp Med, vol. 70, no. 2, pp. 170–175, Apr. 2020, doi: 10.30802/AALAS-CM-19-000022.

[33] B. Boonsri, K. Buddhachat, V. Punyapornwithaya, M. Phatsara, and K. Nganvongpanit, “Determination of whether morphometric analysis of vertebrae in the domestic cat (Felis catus) is related to sex or skull shape,” Anat Sci Int, vol. 95, no. 3, pp. 387–398, Jun. 2020, doi: 10.1007/s12565-020-00533-3.

[34] L. A. Geddes and L. E. Baker, “THE SPECIFIC RESISTANCE OF BIOLOGICAL MATERIAL--A COMPENDIUM OF DATA FOR THE BIOMEDICAL ENGINEER AND PHYSIOLOGIST*t,” p. 23.

[35] J. B. Ranck Jr. and S. L. BeMent, “The specific impedance of the dorsal columns of cat: An anisotropic medium,” Experimental Neurology, vol. 11, no. 4, pp. 451–463, Apr. 1965, doi: 10.1016/0014-4886(65)90059-2.

[36] S. J. Nagel et al., “Spinal dura mater: biophysical characteristics relevant to medical device development,” Journal of Medical Engineering & Technology, vol. 42, no. 2, pp. 128–139, Feb. 2018, doi: 10.1080/03091902.2018.1435745.

[37] “Dielectric Properties » IT’IS Foundation.” https://itis.swiss/virtual-population/tissue-properties/database/dielectric-properties/ (accessed Jul. 08, 2022).

[38] “Polyimide.” http://www.mit.edu/~6.777/matprops/polyimide.htm (accessed Apr. 30, 2021).

[39] R. D. Graham, T. M. Bruns, B. Duan, and S. F. Lempka, “The Effect of Clinically Controllable Factors on Neural Activation During Dorsal Root Ganglion Stimulation,” Neuromodulation: Technology at the Neural Interface, vol. 24, no. 4, pp. 655–671, 2021, doi: 10.1111/ner.13211.

[40] “Rhinoceros®.” Robert McNeel & Associates, Seattle, WA.

[41] C. C. McIntyre, A. G. Richardson, and W. M. Grill, “Modeling the Excitability of Mammalian Nerve Fibers: Influence of Afterpotentials on the Recovery Cycle,” Journal of Neurophysiology, vol. 87, no. 2, pp. 995–1006, Feb. 2002, doi: 10.1152/jn.00353.2001.

[42] K. H. Lee, K. Chung, J. M. Chung, and R. E. Coggeshall, “Correlation of cell body size, axon size, and signal conduction velocity for individually labelled dorsal root ganglion cells in the cat,” Journal of Comparative Neurology, vol. 243, no. 3, pp. 335–346, 1986, doi: 10.1002/cne.902430305.

[43] R. Amir and M. Devor, “Electrical Excitability of the Soma of Sensory Neurons Is Required for Spike Invasion of the Soma, but Not for Through-Conduction,” Biophysical Journal, vol. 84, no. 4, pp. 2181–2191, Apr. 2003, doi: 10.1016/S0006-3495(03)75024-3.

[44] M. Ito and I. Takahashi, “Impulse conduction through spinal ganglion,” in Electrical activity of single cells, 1st ed. Igaku Shoin, Tokyo, 1960, pp. 159–179.

[45] Y. Zheng, P. Liu, L. Bai, J. S. Trimmer, B. P. Bean, and D. D. Ginty, “Deep Sequencing of Somatosensory Neurons Reveals Molecular Determinants of Intrinsic Physiological Properties,” Neuron, vol. 103, no. 4, pp. 598–616.e7, Aug. 2019, doi: 10.1016/j.neuron.2019.05.039.

[46] Z. F. Mainen and T. J. Sejnowski, “Influence of dendritic structure on firing pattern in model neocortical neurons,” Nature, vol. 382, no. 6589, Art. no. 6589, Jul. 1996, doi: 10.1038/382363a0.

[47] P. L. Sheets, J. O. Jackson, S. G. Waxman, S. D. Dib-Hajj, and T. R. Cummins, “A Nav1.7 channel mutation associated with hereditary erythromelalgia contributes to neuronal hyperexcitability and displays reduced lidocaine sensitivity,” The Journal of Physiology, vol. 581, no. 3, pp. 1019–1031, 2007, doi: 10.1113/jphysiol.2006.127027.

[48] J. L. Gaines, K. E. Finn, J. P. Slopsema, L. A. Heyboer, and K. H. Polasek, “A model of motor and sensory axon activation in the median nerve using surface electrical stimulation,” J Comput Neurosci, vol. 45, no. 1, pp. 29–43, Aug. 2018, doi: 10.1007/s10827-018-0689-5.

[49] J. Goodman and J. Weare, “Ensemble samplers with affine invariance,” Communications in Applied Mathematics and Computational Science, vol. 5, no. 1, pp. 65–80, Jan. 2010, doi: 10.2140/camcos.2010.5.65.

[50] D. Foreman-Mackey, D. W. Hogg, D. Lang, and J. Goodman, “emcee: The MCMC Hammer,” PASP, vol. 125, no. 925, p. 306, Feb. 2013, doi: 10.1086/670067.

[51] C. Adams, W. Stroberg, R. A. DeFazio, S. Schnell, and S. M. Moenter, “Gonadotropin-Releasing Hormone (GnRH) Neuron Excitability Is Regulated by Estradiol Feedback and Kisspeptin,” J. Neurosci., vol. 38, no. 5, pp. 1249–1263, Jan. 2018, doi: 10.1523/JNEUROSCI.2988-17.2017.

[52] H. Lee et al., “Molecular Determinants of Mechanical Itch Sensitization in Chronic Itch,” Frontiers in Molecular Neuroscience, Jun. 2022, doi: 10.3389/fnmol.2022.937890.

[53] E. Matsumoto and J. Rosenbluth, “Plasma membrane structure at the axon hillock, initial segment and cell body of frog dorsal root ganglion cells,” J Neurocytol, vol. 14, no. 5, pp. 731–747, Oct. 1985, doi: 10.1007/BF01170825.

[54] R. D. Graham, T. M. Bruns, B. Duan, and S. F. Lempka, “Dorsal root ganglion stimulation for chronic pain modulates Aβ-fiber activity but not C-fiber activity: A computational modeling study,” Clinical Neurophysiology, vol. 130, no. 6, pp. 941–951, Jun. 2019, doi: 10.1016/j.clinph.2019.02.016.

[55] T. Fukuoka, K. Kobayashi, H. Yamanaka, K. Obata, Y. Dai, and K. Noguchi, “Comparative study of the distribution of the α-subunits of voltage-gated sodium channels in normal and axotomized rat dorsal root ganglion neurons,” Journal of Comparative Neurology, vol. 510, no. 2, pp. 188–206, 2008, doi: 10.1002/cne.21786.

[56] C. Ho and M. E. O’Leary, “Single-cell analysis of sodium channel expression in dorsal root ganglion neurons,” Molecular and Cellular Neuroscience, vol. 46, no. 1, pp. 159–166, Jan. 2011, doi: 10.1016/j.mcn.2010.08.017.

[57] M. N. Rasband, E. W. Park, T. W. Vanderah, J. Lai, F. Porreca, and J. S. Trimmer, “Distinct potassium channels on pain-sensing neurons,” PNAS, vol. 98, no. 23, pp. 13373–13378, Nov. 2001, doi: 10.1073/pnas.231376298.

[58] M. S. Gold, M. J. Shuster, and J. D. Levine, “Characterization of six voltage-gated K+ currents in adult rat sensory neurons,” Journal of Neurophysiology, vol. 75, no. 6, pp. 2629–2646, Jun. 1996, doi: 10.1152/jn.1996.75.6.2629.

[59] A. C. Jackson and B. P. Bean, “State-Dependent Enhancement of Subthreshold A-Type Potassium Current by 4-Aminopyridine in Tuberomammillary Nucleus Neurons,” Journal of Neuroscience, vol. 27, no. 40, pp. 10785–10796, Oct. 2007, doi: 10.1523/JNEUROSCI.0935-07.2007.

[60] H. R. Koerber, R. E. Druzinsky, and L. M. Mendell, “Properties of somata of spinal dorsal root ganglion cells differ according to peripheral receptor innervated,” Journal of Neurophysiology, vol. 60, no. 5, pp. 1584–1596, Nov. 1988, doi: 10.1152/jn.1988.60.5.1584.

[61] L. Djouhri, L. Bleazard, and S. N. Lawson, “Association of somatic action potential shape with sensory receptive properties in guinea-pig dorsal root ganglion neurones,” The Journal of Physiology, vol. 513, no. 3, pp. 857–872, 1998, doi: 10.1111/j.1469-7793.1998.857ba.x.

[62] P. J. Waddell and S. N. Lawson, “Electrophysiological properties of subpopulations of rat dorsal root ganglion neurons in vitro,” Neuroscience, vol. 36, no. 3, pp. 811–822, Jan. 1990, doi: 10.1016/0306-4522(90)90024-X.

[63] G. R. Holt and C. Koch, “Electrical Interactions via the Extracellular Potential Near Cell Bodies,” p. 16.

[64] A. R. Shifman and J. E. Lewis, “ELFENN: A Generalized Platform for Modeling Ephaptic Coupling in Spiking Neuron Models,” Front. Neuroinform., vol. 13, 2019, doi: 10.3389/fninf.2019.00035.

[65] C. A. Anastassiou, R. Perin, H. Markram, and C. Koch, “Ephaptic coupling of cortical neurons,” Nat Neurosci, vol. 14, no. 2, pp. 217–223, Feb. 2011, doi: 10.1038/nn.2727.

[66] R. Plonsey and R. C. Barr, Bioelectricity: a quantitative approach, 3rd ed. New York, NY: Springer, 2007.

[67] P. Virtanen et al., “SciPy 1.0: fundamental algorithms for scientific computing in Python,” Nat Methods, vol. 17, no. 3, pp. 261–272, Mar. 2020, doi: 10.1038/s41592-019-0686-2.

[68] “Grapevine User Manual.” Ripple, LLC, Dec. 2016.

[69] C. Luscher, J. Streit, R. Quadroni, and H. R. Luscher, “Action potential propagation through embryonic dorsal root ganglion cells in culture. I. Influence of the cell morphology on propagation properties,” Journal of Neurophysiology, vol. 72, no. 2, pp. 622–633, Aug. 1994, doi: 10.1152/jn.1994.72.2.622.

[70] D. Sundt, N. Gamper, and D. B. Jaffe, “Spike propagation through the dorsal root ganglia in an unmyelinated sensory neuron: a modeling study,” Journal of Neurophysiology, vol. 114, no. 6, pp. 3140–3153, Sep. 2015, doi: 10.1152/jn.00226.2015.

[71] C. Gold, D. A. Henze, C. Koch, and G. Buzsáki, “On the Origin of the Extracellular Action Potential Waveform: A Modeling Study,” Journal of Neurophysiology, vol. 95, no. 5, pp. 3113–3128, May 2006, doi: 10.1152/jn.00979.2005.

[72] D. A. Henze, Z. Borhegyi, J. Csicsvari, A. Mamiya, K. D. Harris, and G. Buzsáki, “Intracellular Features Predicted by Extracellular Recordings in the Hippocampus In Vivo,” Journal of Neurophysiology, vol. 84, no. 1, pp. 390–400, Jul. 2000, doi: 10.1152/jn.2000.84.1.390.

[73] M. J. Stebbing, S. Eschenfelder, H.-J. Häbler, M. C. Acosta, W. Jänig, and E. M. McLachlan, “Changes in the action potential in sensory neurones after peripheral axotomy in vivo,” NeuroReport, vol. 10, no. 2, pp. 201–206, Feb. 1999.

[74] S. Matsuda et al., “Phylogenetic investigation of Dogiel’s pericellular nests and Cajal’s initial glomeruli in the dorsal root ganglion,” Journal of Comparative Neurology, vol. 491, no. 3, pp. 234–245, 2005, doi: 10.1002/cne.20713.

[75] A. A. Jiman et al., “Multi-channel intraneural vagus nerve recordings with a novel high-density carbon fiber microelectrode array,” Sci Rep, vol. 10, no. 1, Art. no. 1, Sep. 2020, doi: 10.1038/s41598-020-72512-7.

[76] J. J. FitzGerald et al., “A regenerative microchannel neural interface for recording from and stimulating peripheral axons in vivo,” J. Neural Eng., vol. 9, no. 1, p. 016010, Jan. 2012, doi: 10.1088/1741-2560/9/1/016010.

[77] T. Guo et al., “Extracellular single-unit recordings from peripheral nerve axons in vitro by a novel multichannel microelectrode array,” Sensors and Actuators B: Chemical, vol. 315, p. 128111, Jul. 2020, doi: 10.1016/j.snb.2020.128111.

[78] D. A. Adamos, E. K. Kosmidis, and G. Theophilidis, “Performance evaluation of PCA-based spike sorting algorithms,” Computer Methods and Programs in Biomedicine, vol. 91, no. 3, pp. 232–244, Sep. 2008, doi: 10.1016/j.cmpb.2008.04.011.

[79] M. Devor, “Unexplained peculiarities of the dorsal root ganglion,” Pain, vol. 82, pp. S27–S35, Aug. 1999, doi: 10.1016/S0304-3959(99)00135-9.

[80] J. Malmivuo and R. Plonsey, Bioelectromagnetism: principles and applications of bioelectric and biomagnetic fields. New York: Oxford University Press, 1995.

[81] A. Tveito et al., “An Evaluation of the Accuracy of Classical Models for Computing the Membrane Potential and Extracellular Potential for Neurons,” Front. Comput. Neurosci., vol. 11, 2017, doi: 10.3389/fncom.2017.00027.

[82] M. Kadwani, R. D. Graham, Z. J. Sperry, S. F. Lempka, and T. M. Bruns, “Computational model of neural recordings in dorsal root ganglia,” presented at the Society for Neuroscience Annual Meeting, Chicago, IL, Oct. 2019.

[83] M. P. Alvarez, M. T. Solas, I. Suarez, and B. Fernandez, “Glial Fibrillary Acidic Protein-Like Immunoreactivity in Cat Satellite Cells of Sympathetic Ganglia,” CTO, vol. 136, no. 1, pp. 9–11, 1989, doi: 10.1159/000146789.

[84] R. S. Nascimento, M. F. Santiago, S. A. Marques, S. Allodi, and A. M. B. Martinez, “Diversity among satellite glial cells in dorsal root ganglia of the rat,” Braz J Med Biol Res, vol. 41, no. 11, pp. 1011–1017, Oct. 2008, doi: 10.1590/S0100-879X2008005000051.

[85] A. R. Harris, “Understanding charge transfer on the clinically used conical Utah electrode array: charge storage capacity, electrochemical impedance spectroscopy and effective electrode area,” J. Neural Eng., vol. 18, no. 2, p. 025001, Feb. 2021, doi: 10.1088/1741-2552/abd897.

[86] K. S. Mathews et al., “Acute Monitoring of Genitourinary Function Using Intrafascicular Electrodes: Selective Pudendal Nerve Activity Corresponding to Bladder Filling, Bladder Fullness, and Genital Stimulation,” Urology, vol. 84, no. 3, pp. 722–729, Sep. 2014, doi: 10.1016/j.urology.2014.05.021.

