## Supplemental for "Multiformity of extracellular microelectrode recordings from Aδ neurons in the dorsal root ganglia: A computational modeling study"

**Manuscript Title:**

**Supplemental Figures**

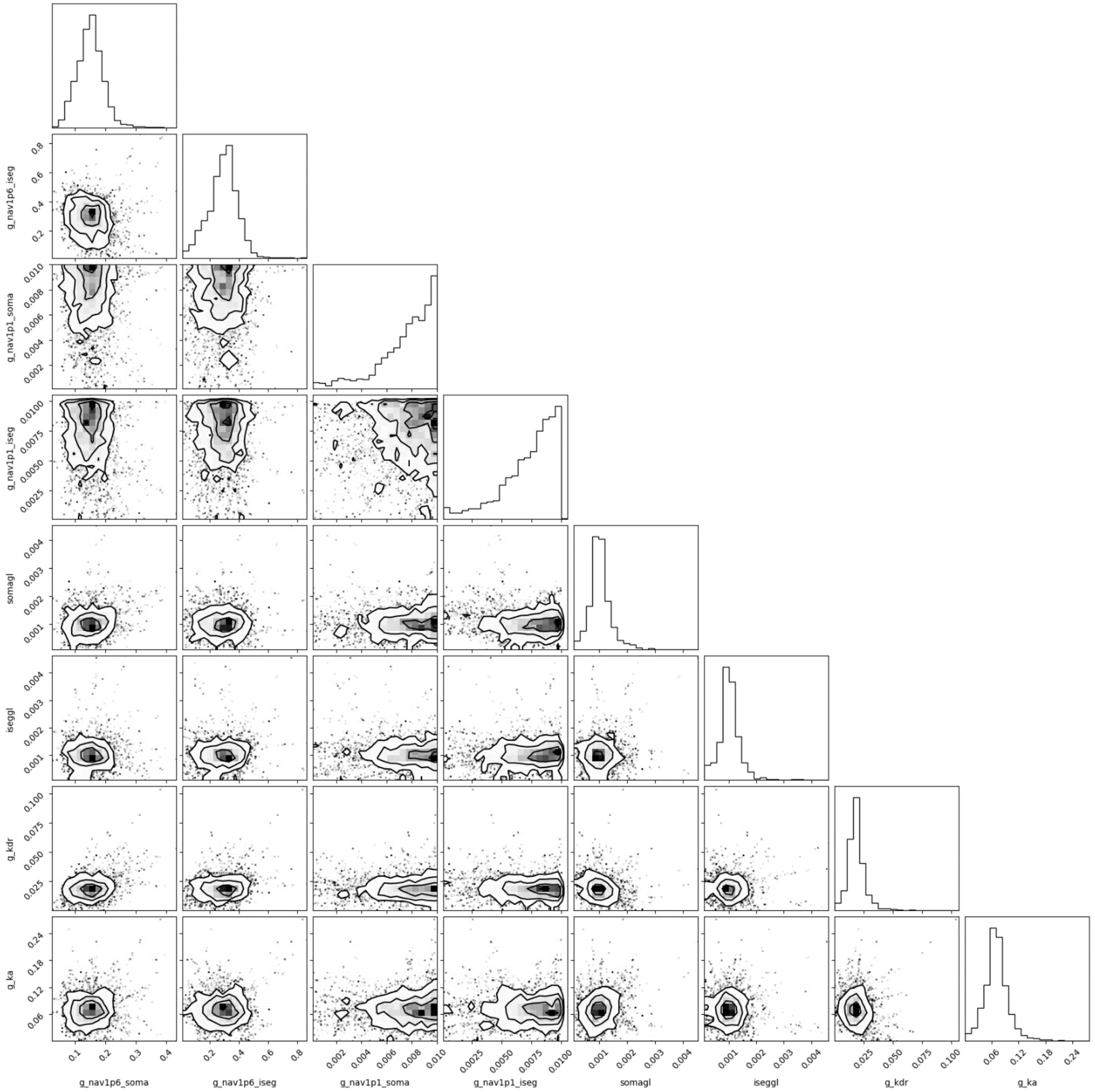


Figure S1: Contour plots and probability distributions generated by the Markov Chain Monte Carlo (MCMC) optimization method. Each dot that makes up the contour plots represents a single iteration of model parameters that were tested. Resulting contour plots show any correlations between parameters. The diagonal shows the probability distribution for the value of each parameter.

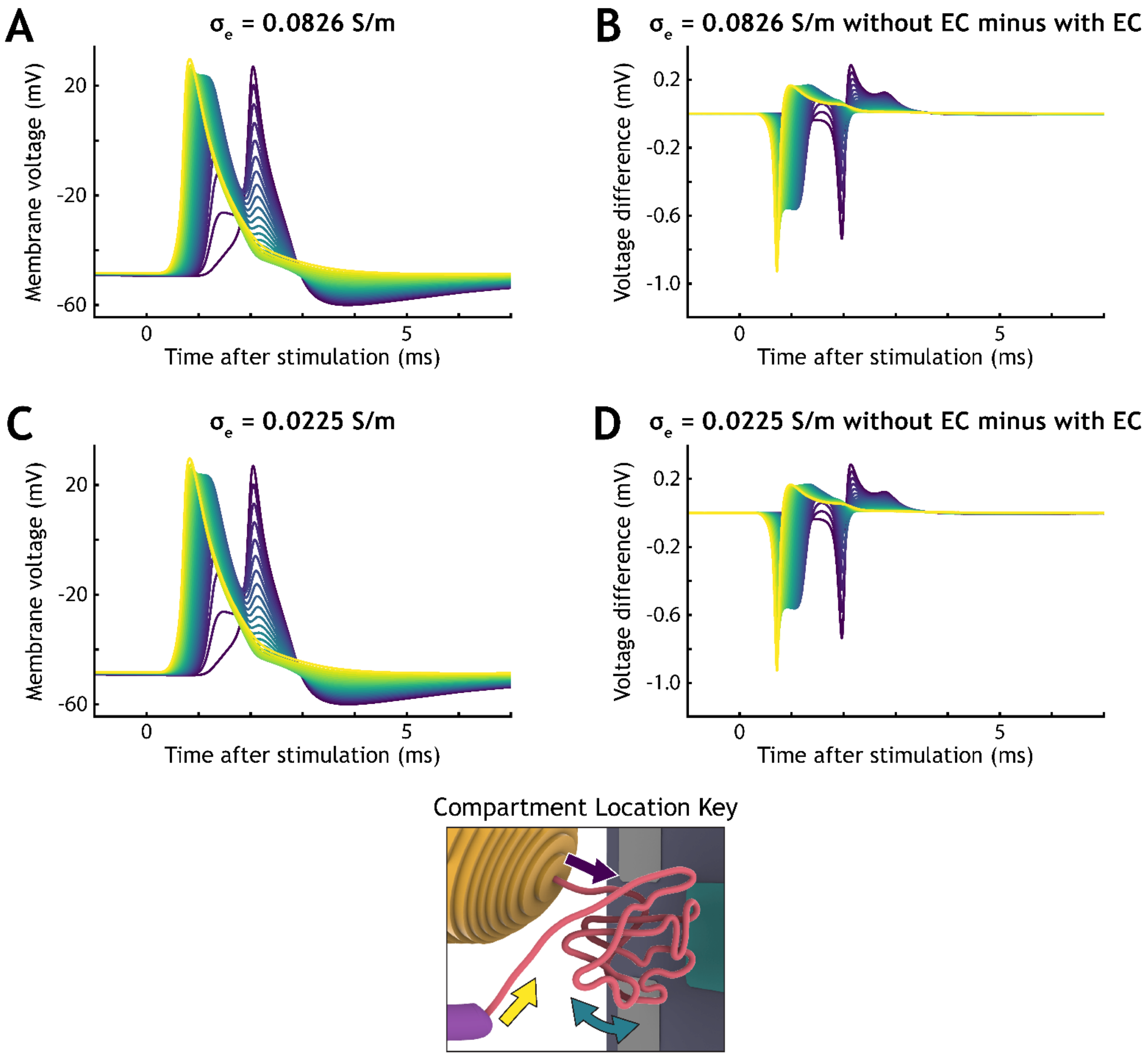


<u>Figure S2:</u> Comparing the glomerular initial segment (GIS) intracellular action potential voltage profile (ICAPVP) from simulations that used auto-ephaptic coupling (EC) to those without. Each line represents the intracellular voltage of a single GIS compartment. The color of the line corresponds with the location along the GIS, as indicated by the Compartment Location Key.

(A) Results from a simulation that implemented auto-ephaptic coupling with an extracellular (i.e., dorsal root ganglion (DRG) grey matter) conductivity of 0.0826 S/m.

(B) Results in (A) subtracted from the results of a simulation that did not implement auto-ephaptic coupling (Figure 4B).

(C) The same as (A), but instead using a conductivity of 0.0225 S/m.

(D) The same as (B), but instead subtracting the results in (C) from the results in Figure 4B.

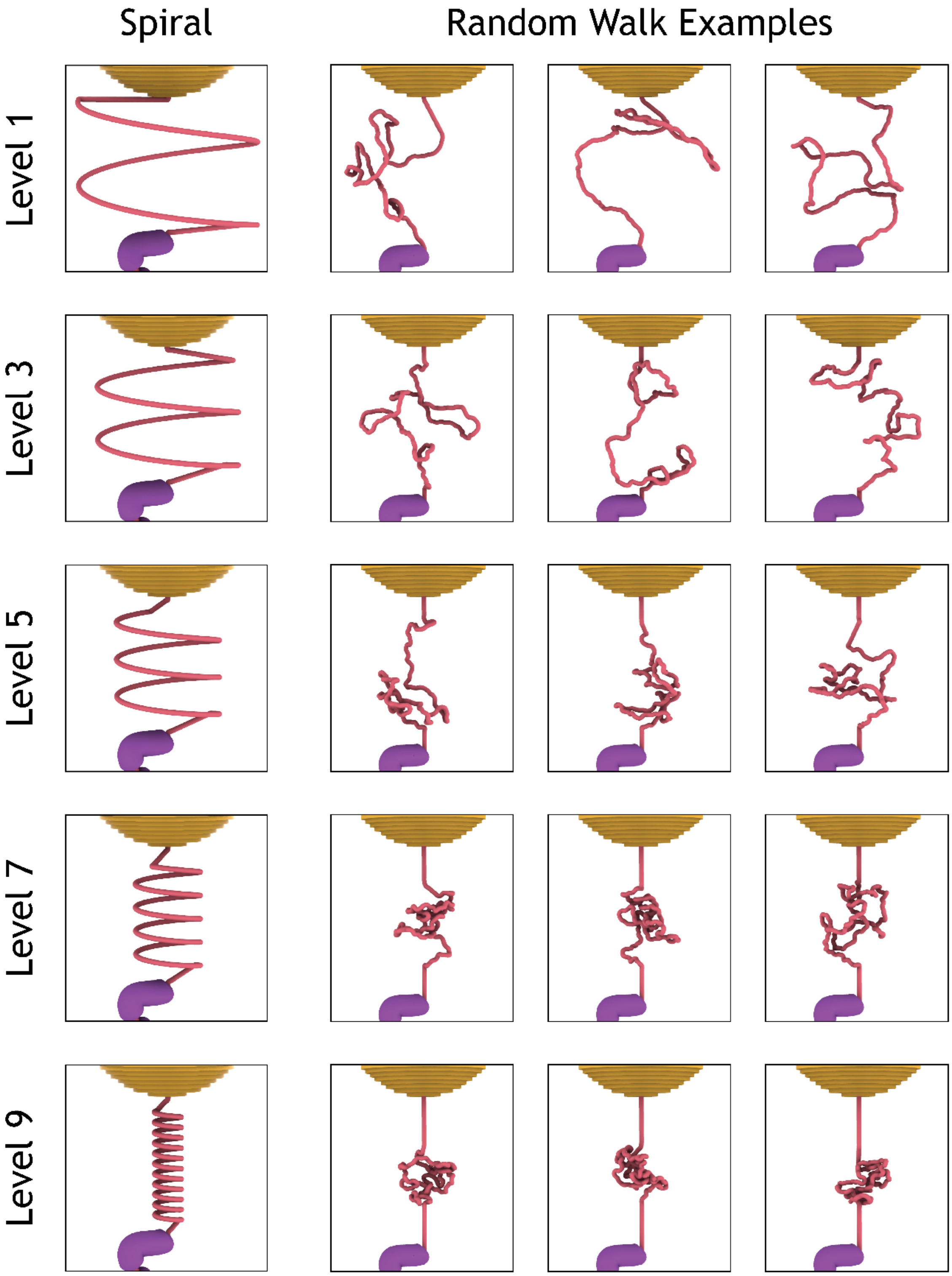


Figure S3: Different GIS trajectories that were used. The first column shows the spiral trajectories for different tortuosity levels. The second through fourth columns show examples of random walk trajectories for different tortuosity levels.

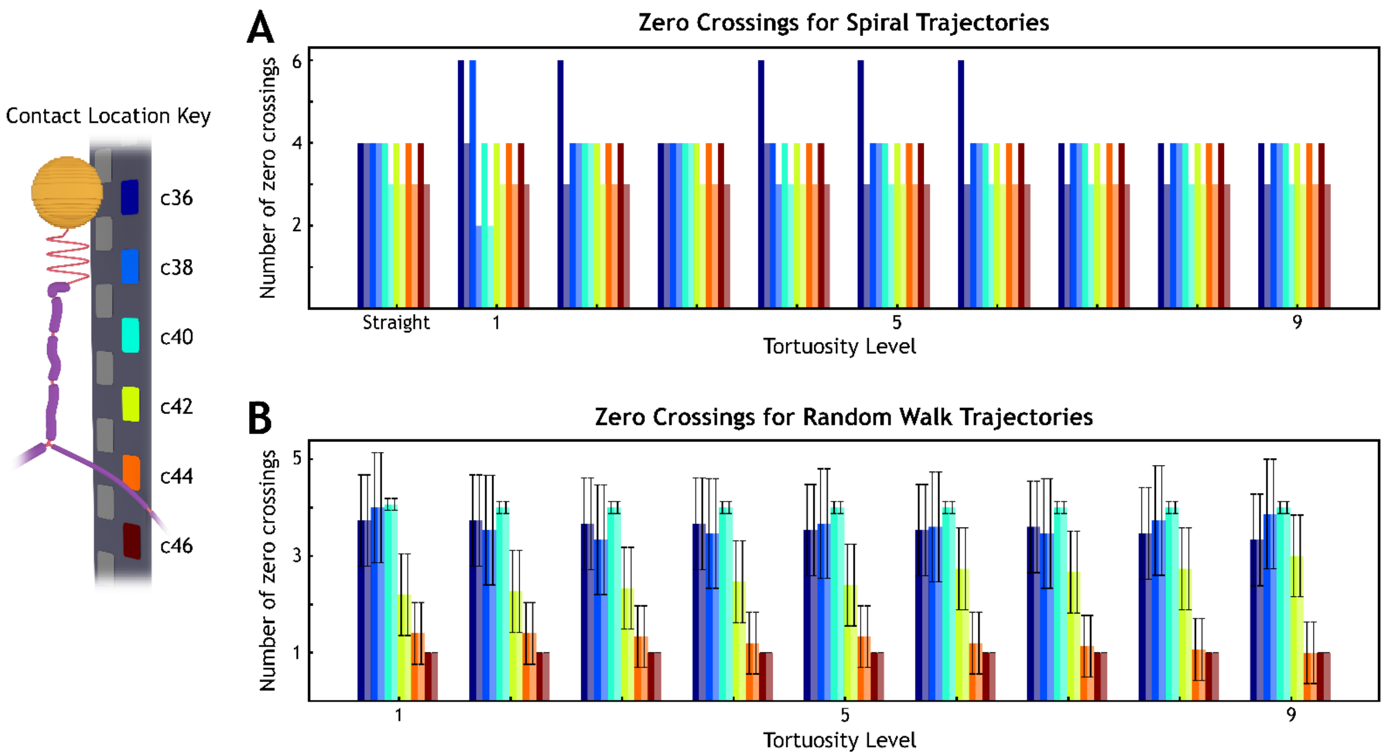


Figure S4: Number of zero crossings across levels of each tortuosity type. The color of the bar corresponds with the color of the electrode contact on the key to the left. Darker bars represent results from simulations that did not implement ephaptic coupling, and lighter bars lines represent results that did.

(A) Number of zero crossings for spiral trajectories.

(B) Number of zero crossings for all 30 random walk trajectories at each tortuosity level.

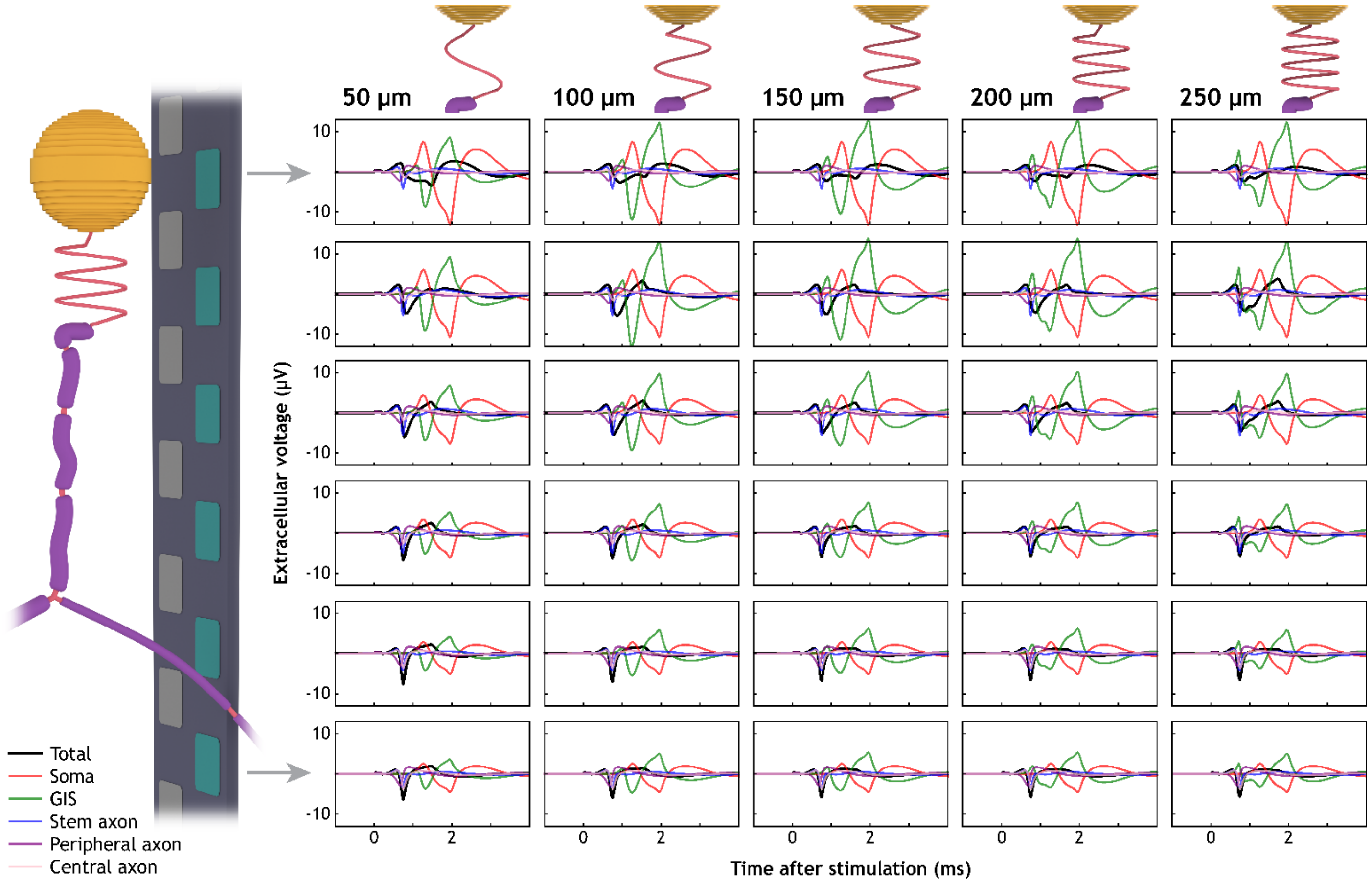


Figure S5: Contributions of different neural components to the extracellular waveforms using an extracellular (DRG grey matter) conductivity of $\sigma_e$ = 0.26 S/m. Each row contains waveforms from the same electrode contact, as indicated by the model rendering on the left. Each column contains waveforms from neurons with the GIS length as indicated by the renderings and titles at the top of the column. The total waveform is shown in bold black, while contributions from separate components are colored via the key on the bottom left.

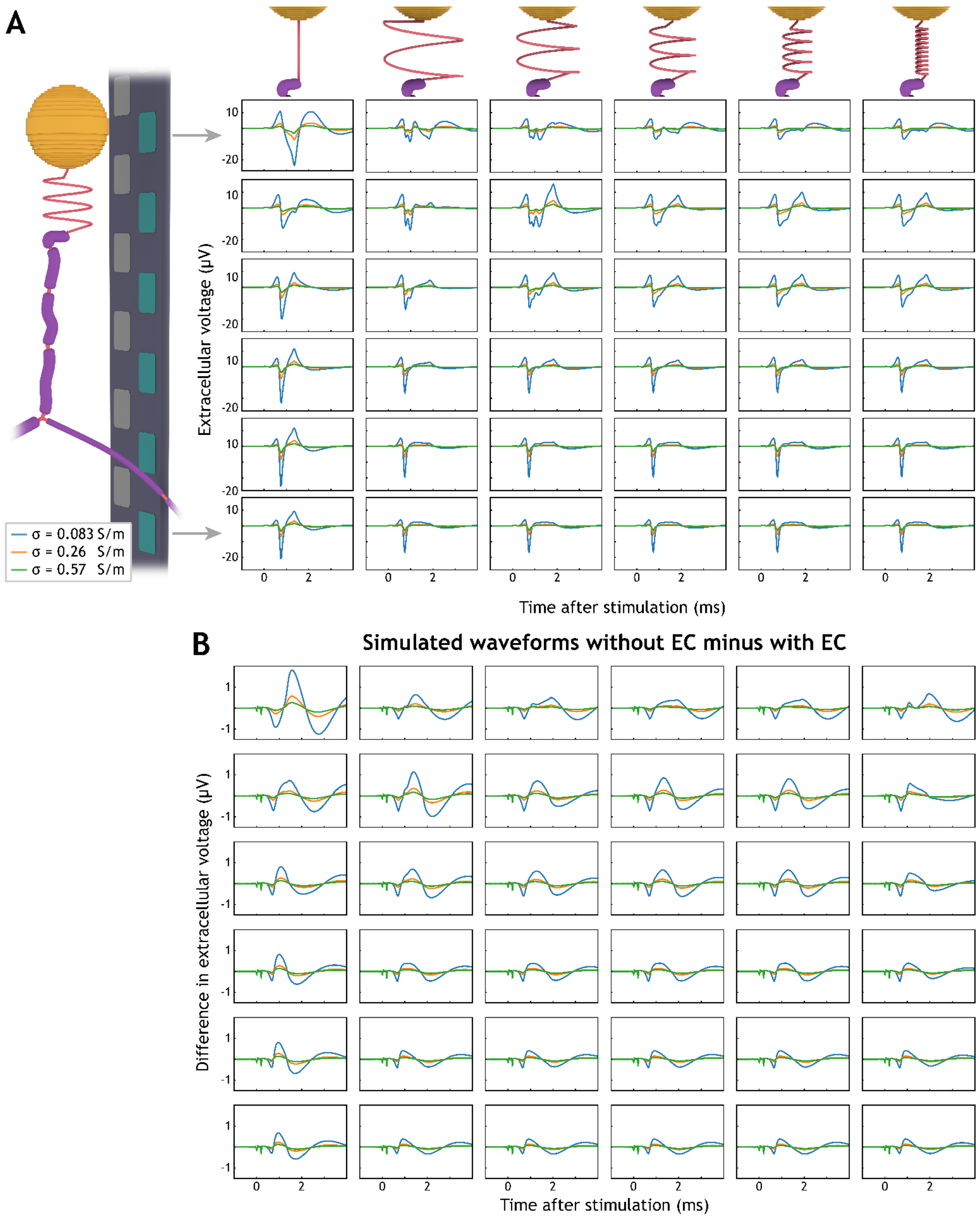


Figure S6: Differences in extracellular waveforms simulated with and without implementing ephaptic coupling. (A) Shows the same simulated results as Figure 5, except these simulations implemented ephaptic coupling. (B) The results in Figure S6A subtracted from the results in Figure 5. The rows, columns, and line color legend are the same as in Figure 5 and Figure S6A.

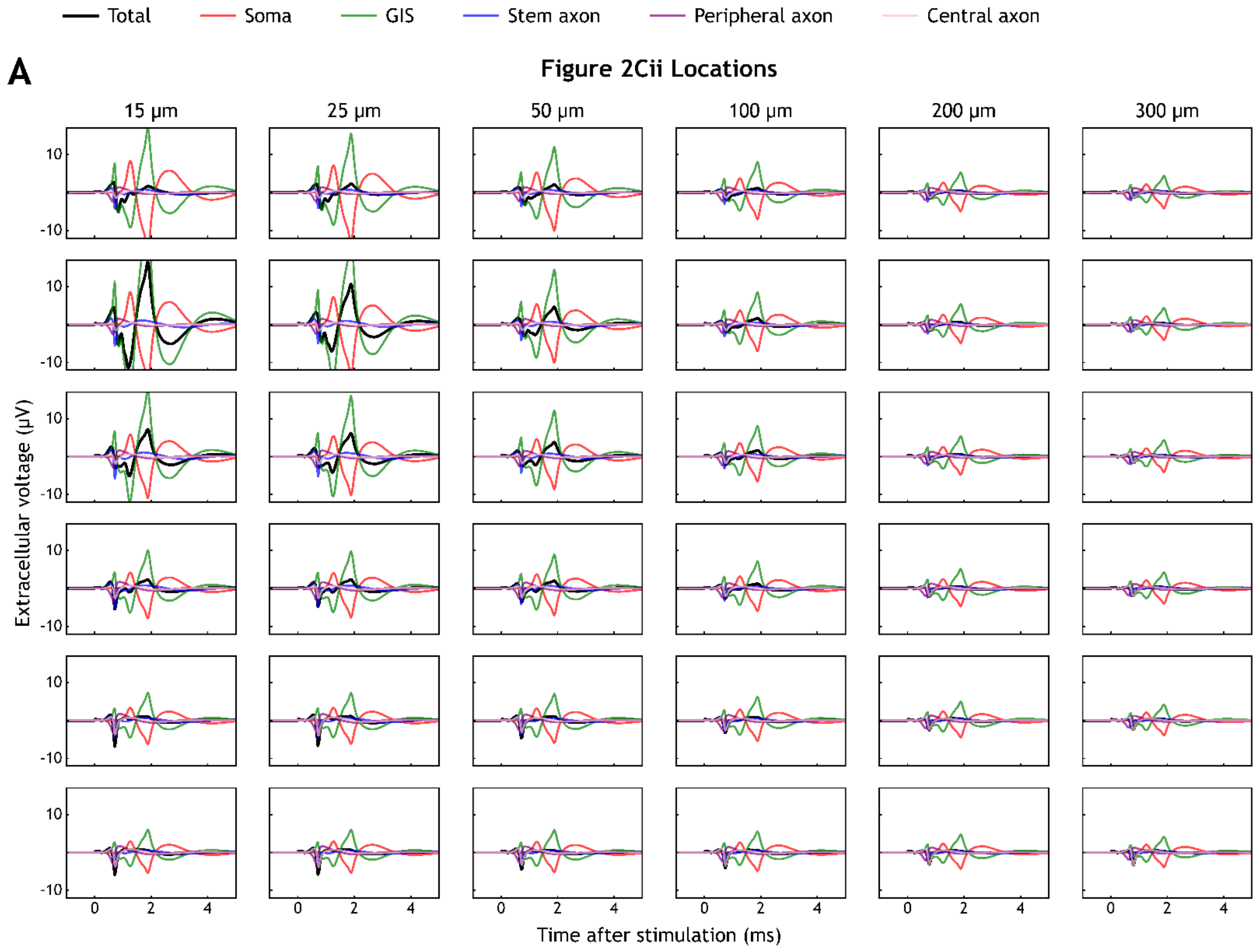

Total
Soma
GIS
Stem axon
Peripheral axon
Central axon
A
Figure 2Cii Locations
15 μm
25 μm
50 μm
100 μm
200 μm
300 μm
10
-10
Extracellular voltage (μV)
0
2
4
Time after stimulation (ms)

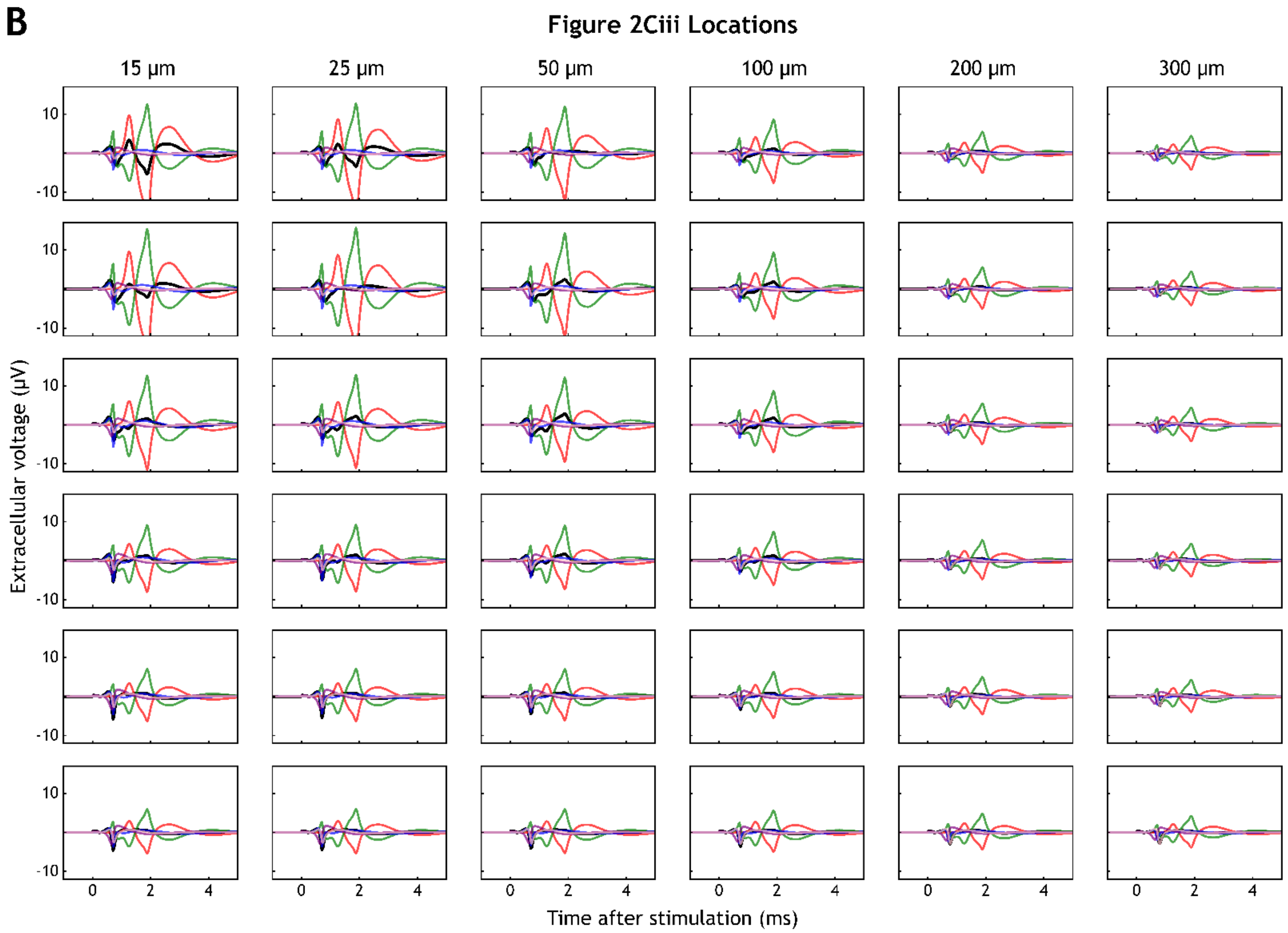


Figure S7: Contributions of different components of the neuron model to the overall waveform, based on microelectrode array distance from the GIS.

(A) The same waveforms shown in Figure 9A, but each row corresponds to the electrode contact shown Figure S6A. Each column corresponds with the distance between the GIS and the electrode contact, as in Figure 9A. The different colored lines correspond with different components of the neuron model, for which the key is at the top of the figure.

(B) The same waveforms shown in shown in Figure 9B, but separated by electrode contact as in (A).

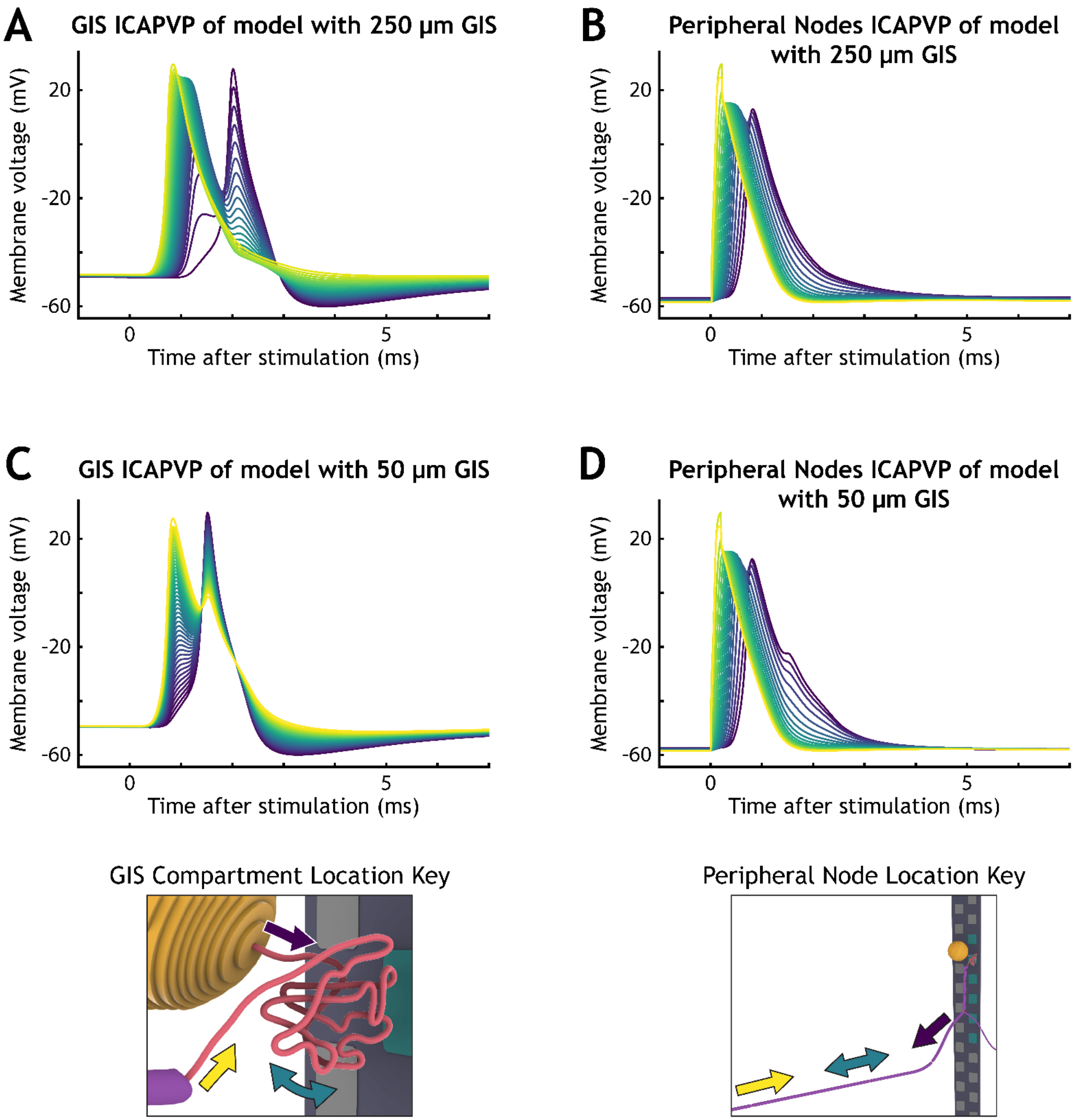


Figure S8: Intracellular voltage profiles of the GIS of two different lengths.
(A) The ICAPVP of GIS compartments at intervals of ten compartments across the 250 compartments in this iteration of the neuron model. The colors of the lines correspond with the location of the compartments, as indicated by the key on the bottom left.
(B) The ICAPVP of peripheral branch node compartments of the iteration of the neuron model with a 250 µm GIS. The colors of the lines correspond with the location of the compartments, as indicated by the key on the bottom right.
(C) The same as (A), but for an iteration of the neuron model with a 50 µm GIS.
(D) The same as (B), but for an iteration of the neuron model with a 50 µm GIS.

**Supplemental Methods**

Random walk pathway algorithm

We used three methods within the algorithm to increase the tortuosity of the paths it generated across the tortuosity levels. First, we delimited a sphere in the region of the GIS to constrain the volume in which the random walk algorithm can explore. Less tortuous paths had larger volumes of space where the trajectory could be built, and more tortuous paths had a smaller volume of space, leading to more compact trajectories between the top of the stem axon and the bottom of the soma. Second, as the tortuosity level increased, the algorithm permitted smaller distances between non-adjacent compartments within the path. Third, more tortuous paths also had an increased range for randomly selected angles between consecutive compartments. We selected the largest and smallest spheres of constraint (for the least and most tortuous paths, respectively) to be similar in overall size to the largest and smallest of the spiral trajectories, so that tortuosity was comparable between the two types of trajectories. Further, we used two tactics within the algorithm to generate random walk trajectories that ended at the same global coordinates, ensuring the soma was constructed in the same location relative to the electrodes, while maintaining a consistent path length. First, for more tortuous trajectories with smaller spheres of path constraint, the trajectories included straight paths below and above the sphere of constraint, which maintained a consistent overall path height. Second, bias for the direction of the next compartment towards the top of the sphere of constraint increased as a trajectory's length was assembled.

To generate one random walk trajectory of a given tortuosity, we generated multiple iterations of random walk trajectories with the algorithm until the random portion of the path ended near the top of the sphere of constraint. Because the random portion path would end near the top of the sphere but effectively never at the exact coordinates, in order to preserve the length of the path, we back-tracked on the trajectory to rebuild the final series of compartments within the sphere. To do this, compartments were incrementally removed from the random portion trajectory and counted until the counted number of compartments could be used to create a straight path towards the top of the sphere of constraint. This would ensure that the next segment(s) of the path, which would be protruding straight from the top of the sphere, always began in the same location. This allowed the overall length of the path to remain consistent with respect to other random walk trajectories generated with the same tortuosity level. Further, we manually checked all random trajectories and paths with hyperacute angles between rebuilt compartments were discarded.

*GitHub® Repository via Zenodo: https://doi.org/10.5281/zenodo.10014936*
